# gamdid: generalized additive models for differential distributions in single cell experiments

**DOI:** 10.64898/2026.06.18.733106

**Authors:** Lucas Beerland, Stijn Vandenbulcke, Christophe Vanderaa, Lennart Martens, Lieven Clement

**Affiliations:** Department of Mathematics, University of Ghent; Department of Biomolecular Medicine, University of Ghent; VIB-UGent Center for Medical Biotechnology; BioOrganic Mass Spectrometry Laboratory (LSMBO), IPHC UMR 7178, University of Strasbourg, CNRS, Strasbourg, France; Infrastructure Nationale de Protéomique ProFI-UAR 2048, Strasbourg, France June 18, 2026

## Abstract

Single-cell proteomics (SCP) generates protein abundance measurements across hundreds to thousands of individual cells, offering unprecedented resolution to study cellular heterogeneity. However, existing differential abundance (DA) methods are limited to detecting shifts in mean expression, leaving biologically relevant differences in shape undetected. Indeed, the specific power of SCP is to identify differences between individual cells in a population, which are typically only found as shape differences rather than in mean expression. We here therefore present gamdid (generalized additive models for differential distributions), a novel statistical framework and R package for differential distribution (DD) analysis in SCP data. gamdid is based on generalized additive models (GAMs) to flexibly model heterogeneous distributions, perform inference and provide interpretable visualizations. Through semi-synthetic benchmarking on two SCP datasets, gamdid demonstrates conservative false discovery rate control and substantially outperforms competing methods for differences in shape, while achieving comparable performance for mean shifts. A spike-in case study further demonstrates the utility of gamdid and its interpretable visualization. Uniquely among DD methods, gamdid supports omnibus testing across more than two groups, with post-hoc pairwise comparisons via stagewise testing, and is specifically tailored for proteomics abundance data.

## Introduction

High-throughput single cell technologies have exponentially improved the resolution at which biological processes can be studied. A key component of such studies is differential abundance (DA) analysis, which aims to identify features that differ between experimental conditions. However, the majority of established DA single-cell methods primarily detect changes in mean abundance and thus overlook other, biologically very informative, differences in distributions, such as changes in variance, skewness, modality, etc., limiting the advantages of single cell over bulk. This limiting focus stems from their origin in bulk omics workflows, where analyses beyond the mean are typically not feasible due to the limited number of available samples. Single-cell datasets, however, provide expression measurements for hundreds to thousands of individual cells, making it possible, and likely necessary, to assess changes in the full distribution of expression values. Indeed, the importance of distribution changes is nicely illustrated in the single-cell proteomics (SCP) data from Leduc et al. [1], which compares melanoma cells with monocytes. Figure 1 visualizes the density of three illustrative preprocessed proteins.

**Figure 1:**
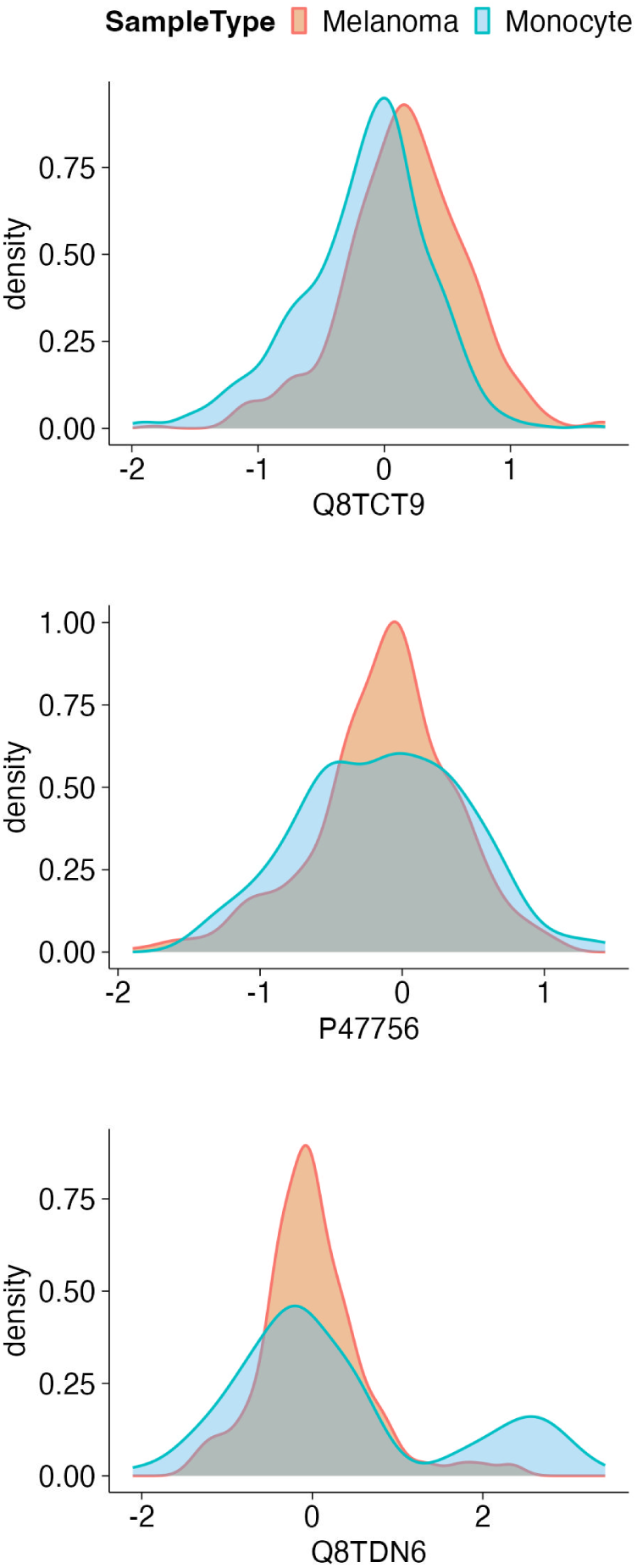
Motivating examples of distributional heterogeneity in single-cell proteomics Density distributions of three proteins from the Leduc single-cell proteomics dataset, comparing melanoma cells (red) and monocytes (blue). The top panel (Q8TCT9) illustrates a conventional mean shift between cell types, which is the type of difference that standard differential abundance (DA) methods are designed to detect. The middle panel (P47756) shows a difference in variance. The bottom panel (Q8TDN6) reveals the emergence of a distinct subpopulation in one cell type, visible as a secondary mode. The latter examples reflect differences in shape rather than mean. These are biologically informative yet systematically missed by conventional DA approaches. They motivate the need for differential distribution (DD) methods capable of detecting the full spectrum of distributional differences in single-cell data.

Conventional DA analysis methods are designed to pick up mean shifts, as seen in the top panel in figure 1, but lack power to prioritize proteins for which the distribution between cell populations mainly differs in shape, as seen in the middle and bottom panels in figure 1. These changes in shape may reflect underlying biological processes that remain undetected with conventional DA methods. For example, the increased spread of the abundance values observed in the middle panel may indicate a heterogeneous cellular response, whereas the pattern in the bottom panel may reflect the emergence of a novel cell state in a subset of cells. To infer such nuanced but relevant effects, more flexible approaches are required, which are referred to as differential distribution (DD) methods.

Several approaches to perform such DD analysis on single cell data already exist, most notably BASiCS [2], scDD [3] and distinct [4]. However, these methods were developed specifically for single-cell RNA-seq data and have neither been tested on, nor tailored for proteomics applications. Moreover, they cannot handle large missing values, a key problem in SCP datasets, and lack interpretability on how the distributions differ. The latter greatly limits the utility of the assessment, as the nature of the difference is an important element in hypothesizing about the possible causes.

Additionally, each approach has methodological limitations: BASiCS is restricted to testing for differences in mean and variance, scDD can only handle count data, and distinct can only be used for two group comparisons.

Therefore we here present gamdid: a novel statistical framework for DD analysis. gamdid is capable of detecting any form of distributional difference between experimental conditions and provides an interpretable visualization of these effects. Importantly, gamdid moreover allows users to localize and visualize the specific regions that are differentially distributed, providing a direct link between statistical inference and visualization. By building on the flexible framework of generalized additive models (GAMs), gamdid thus extends well beyond traditional mean-based comparisons while remaining intuitive and highly interpretable.

## Results

In this section, we first describe the statistical model and inference procedure underlying gamdid. We then evaluate gamdid through a simulation study and a case study, both based on real SCP data. The simulation study benchmarks gamdid across five distributional scenarios. Subsequently, we demonstrate the practical utility of gamdid through a controlled spike-in case study.

### Statistical model and inference

A key advantage of SCP analysis over bulk proteomics is its capacity to assess single-cell heterogeneity. This can result in differences in abundance distributions that are more subtle than mean abundance differences, and are thus overlooked by traditional, bulk-oriented differential analysis methods for proteomics data. Instead of modelling the abundance data with a normal distribution, we here adopt a flexible non-parametric approach to estimate the distribution of the protein abundances in single cells in each group by building upon Lindsay’s method [5]. Specifically, for each feature the density estimation problem is recast into Poisson regression by discretizing the sample space into bins. In other words, we convert the intensities into histogram bin data, see figure 2 panel A, and the bin counts are then considered independent Poisson-distributed observations.

**Figure 2:**
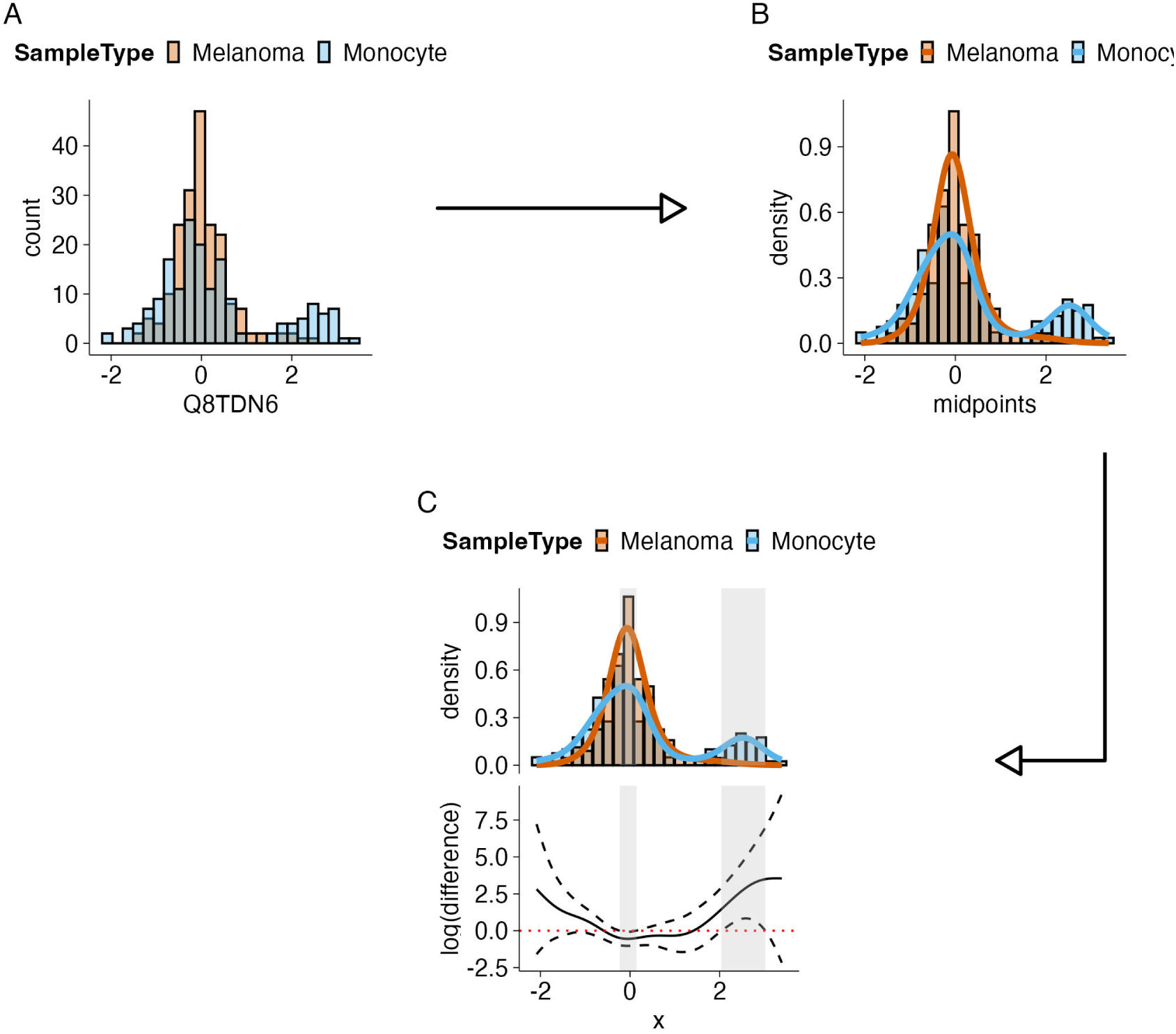
Overview of the gamdid workflow The complete workflow for fitting GAMs to compare distributions is illustrated using a ribosome biogenesis protein from the Leduc dataset. **Panel A**: Raw protein abundance values are discretized into histogram bin counts, recasting the density estimation problem as Poisson regression. This representation preserves distributional shape information that is lost in a mean-based summary. **Panel B**: A Poisson generalized additive model (GAM) is fitted to the bin counts as a function of bin midpoints, allowing each group (melanoma and monocyte) to have its own flexible, non-parametric density curve. The GAM framework accommodates any distributional shape without parametric assumptions. **Panel C (top)**: The fitted density curves generated by gamdid’s visFit function, with grey shading highlighting regions where the distributions differ significantly between groups. **Panel C (bottom)**: The difference smoother representing the log-ratio of densities between groups along with its simultaneous confidence band. Regions where this band excludes zero identify statistically significant local distributional differences, linking inference to an interpretable visualization. This combination of flexible modelling, hypothesis testing, and interpretable visualization is an innovative feature unique to gamdid.

This representation allows for density estimation through Poisson generalized additive models, where expected counts are modeled in function of bin midpoints and by specifying a group specific spline, each group has its own unique density shape. The GAM fits are displayed in figure 2 panel B.

Inference is subsequently carried out using the GAM framework, with differences between groups assessed using Wald-type tests by comparing group-specific model parameters. This yields a p-value for inferring any type of distributional difference both in location and in shape, referred to as the GAM p-value in the remainder of the manuscript.

To increase the power to detect distributional changes that are strongly driven by differences in the mean, the p-value obtained from the GAM is combined with a p-value derived from the msqrob2 framework, a conventional mean-based approach for differential abundance (DA) analysis [6]. The two p-values are integrated using the Harmonic Mean P-value (HMP) method [7]. This results in a final aggregated p-value for each feature, providing a single, robust measure of evidence for any distributional change.

Interestingly, gamdid also provides an interpretable visualization that is directly linked to the inference, by constructing a difference smoother along with a simultaneous confidence band. Regions where this band does not include zero indicate a statistically significant local distributional difference, as illustrated in figure 2 panel C.

### Simulation study for benchmarking

To benchmark gamdid for single cell proteomics applications, we constructed semi-synthetic simulations based on two publicly available mass spectrometry (MS)-based single-cell proteomics (SCP) data sets obtained from the R/Bioconductor package scpdata [8]: the TMT-labeled single-cell Leduc dataset [1] consisting of 1531 cells and 6706 identified peptides, and the label-free single-cell Petrosius dataset [9] consisting of 564 cells and 5729 identified peptides.

For each dataset, preprocessed (see methods section) protein-level summarized assays from one cell type were selected and the corresponding cells were randomly assigned into two mock groups (A and B). Hence, all proteins are non-DD between A and B.

Only proteins with at least 50 non-missing observations per group were retained for benchmarking, consistent with the default setting in the gamdid package. This threshold ensures a sufficient number of observations to reliably characterize the distribution of each feature, as fewer observations may yield an incomplete or misleading representation of the underlying density.

### Simulation of distributional differences

Following the framework established by Korthauer et al. (2016) [3], we introduced five categories of distributional differences by altering protein abundances in cells of group B: differential expression (DE) (a pure mean shift, which corresponds to DA), differential variability (DV), differential component means (DB), differential modality (DM), and differential proportion (DP), as illustrated in Figure 3. These effects were introduced in 20% of randomly selected features by systematic shifting of the abundances in subsets of the cells in group B (for the DE, DB, DM and DP scenarios) or by adding controlled random noise (for the DV scenario).

**Figure 3:**
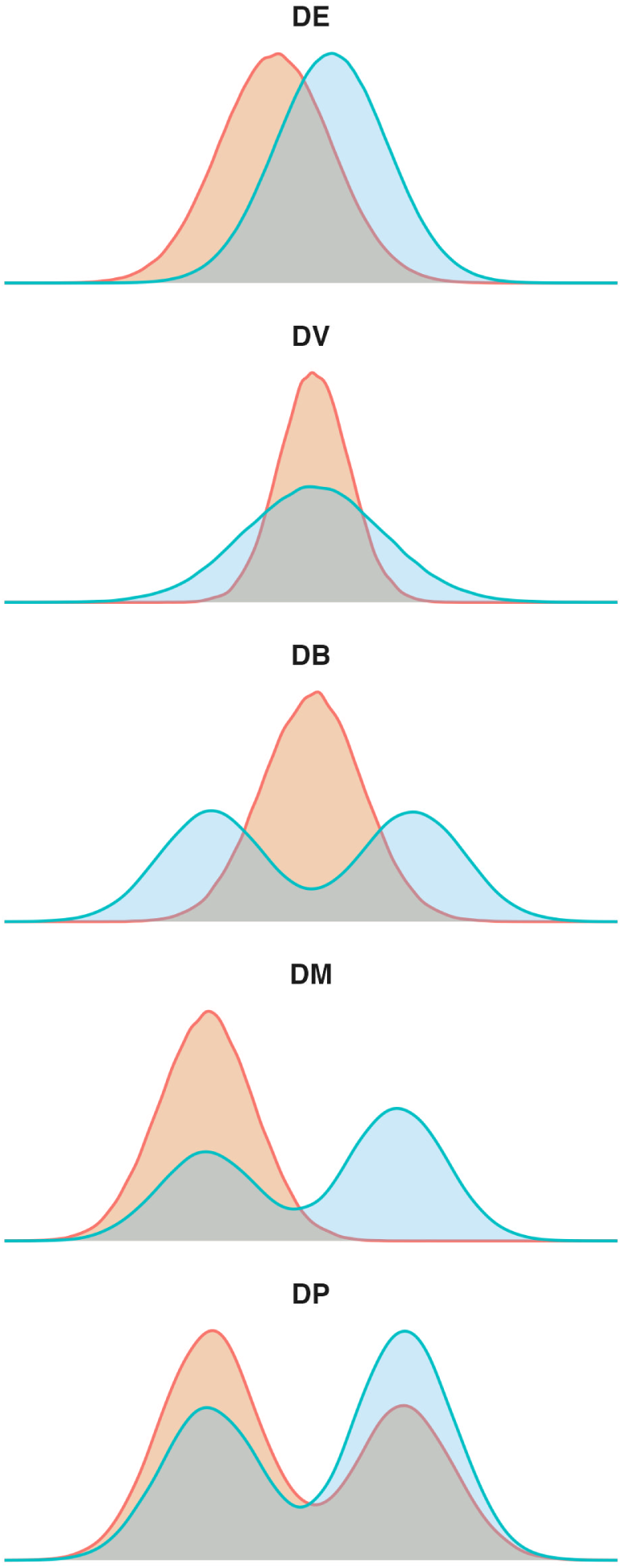
Five simulation scenarios used to benchmark gamdid To evaluate gamdid’s ability to detect different types of distributional differences, five categories of signal, established by Korthauer et al. (2016) [3], were introduced into real single-cell proteomics data. DE (differential expression): a pure shift in the mean, equivalent to conventional differential abundance (DA). DV (differential variability): an increase in variance without a location shift, reflecting e.g. heterogeneous cellular responses. DB (differential component means): a bimodal distribution induced in one group by shifting subsets of cells in opposite directions, without altering the overall mean. DM (differential modality): an asymmetric shift in one tail, creating a difference in skewness or the emergence of a subpopulation. DP (differential proportion): a change in the relative weight of a subpopulation shared between groups.

This semi-synthetic approach leverages real SCP data rather than fully synthetic ”clean“ simulations, and thus preserves realistic distributional characteristics inherent to single-cell proteomics measurements.

### Method comparison

We compare gamdid to distinct as the primary benchmark method. BASiCS and scDD were excluded from the comparison as BASiCS is limited to detecting shifts in mean and variance only, and can therefore not be applied to the majority of the simulations, while scDD requires count data as input, which is incompatible with the continuous abundance measurements obtained in proteomics data. distinct performs group comparisons by computing the average difference between empirical cumulative distribution functions (ECDFs) and can optionally account for subgroup structure in multisample multicell designs. As our datasets, similar to most public SCP datasets, do not have these multisample multicell designs, distinct was applied without this option.

Note that the current implementation of gamdid does not account for subgroup structure.

Method performance is evaluated using false discovery proportion — true positive proportion (FDP-TPP) curves. For a given significance threshold, the FDP represents the proportion of false positives among all features called significant, while the TPP represents the proportion of true positives recovered out of all truly differential features. An FDP-TPP curve allows to assess the power of a method (TPP) at any given level of false discoveries (FDP). This provides a more complete picture of method performance than a single summary statistic, and allows for a direct visual comparison of the trade-off between sensitivity and false discovery control across competing methods. A method can be considered more powerful when the area under its FDP-TPP curve is larger. The dashed vertical line at FDP = 0.05 serves as a reference for the nominal FDR threshold.

### GAM evaluation

We first evaluate the performance of the GAM methodology in isolation (without p-value aggregation via msqrob2) against distinct. The FDP-TPP curves are presented in figures 4 and 5.

**Figure 4:**
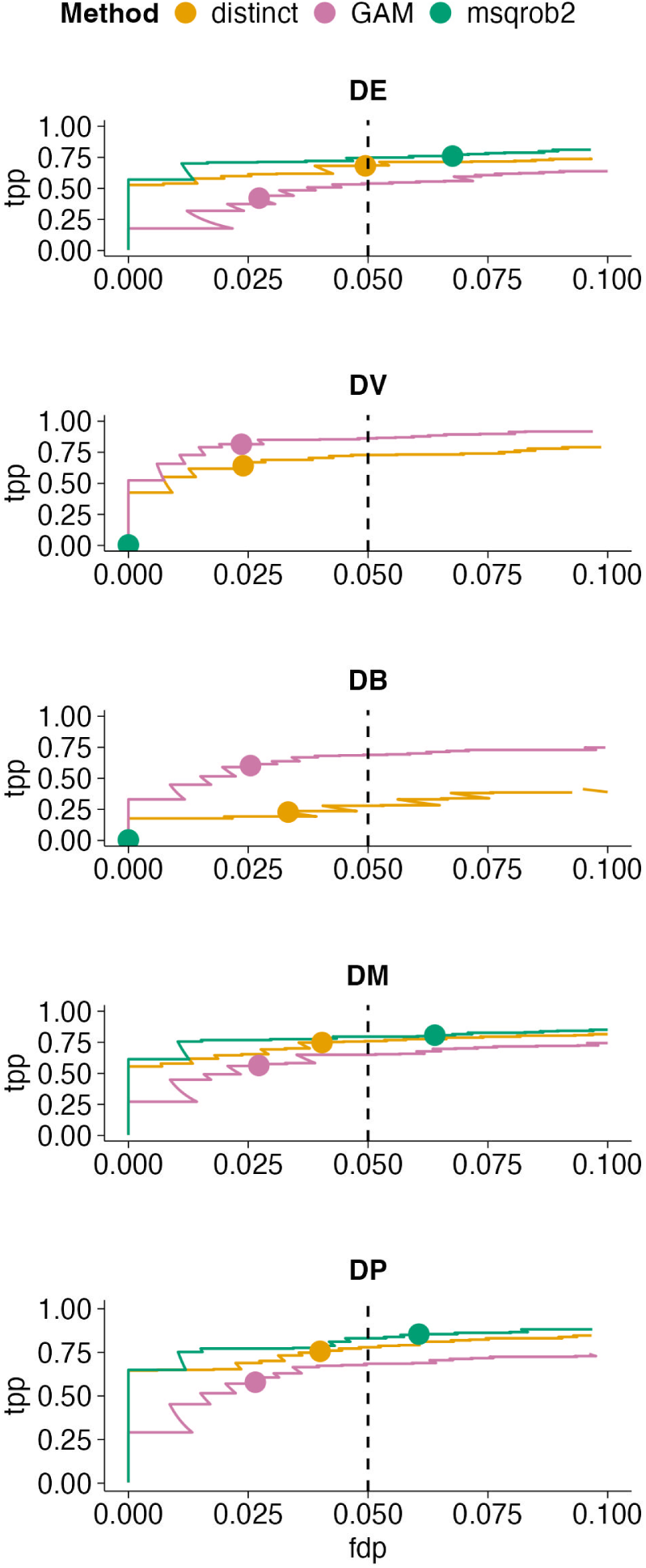
Performance of the standalone GAM method on the Leduc dataset FDP-TPP curves comparing the standalone GAM method, msqrob2, and distinct across the five simulation scenarios in the Leduc dataset (1531 cells, 6706 peptides). Each curve traces the trade-off between true positive proportion (TPP, sensitivity) and false discovery proportion (FDP) as the significance threshold is varied; the dashed vertical line marks FDP = 0.05. The GAM method achieves conservative FDR control across all scenarios. For distributional changes that do not involve a mean shift (DV, DB), the GAM method substantially outperforms distinct, demonstrating its superior sensitivity to shape differences. However, for scenarios where mean shifts contribute to the distributional change (DE, DM, DP), distinct and msqrob2 outperform the standalone GAM method. This complementarity motivates the aggregation of GAM p-values with msqrob2 p-values in the gamdid pipeline.

**Figure 5:**
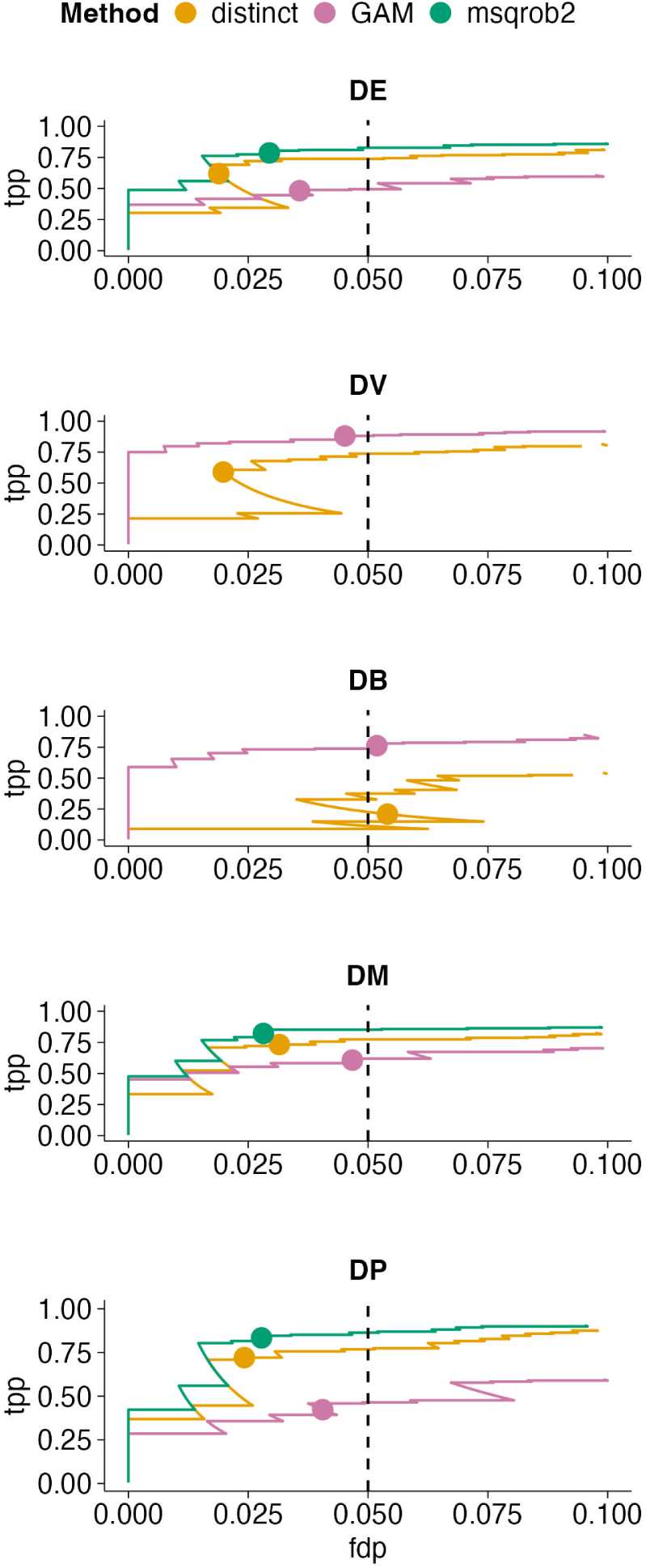
Performance of the standalone GAM method on the Petrosius dataset FDP-TPP curves for the standalone GAM method, msqrob2, and distinct on the Petrosius dataset (564 cells, 5729 peptides), mirroring the analysis in figure 4. The same pattern holds: the GAM method is superior for shape-only changes (DV, DB), while msqrob2 and distinct perform better for DE and partially for DM and DP. Notably, improvements of the GAM method over distinct are more pronounced in the Petrosius dataset than in Leduc, which is attributable to the smaller sample size: contrary to gamdid, distinct relies on permutation-based inference and suffers from reduced null distribution resolution with fewer observations. These results reinforce the case for combining GAM and msqrob2 p-values to obtain a method that is sensitive across all distributional scenarios.

The GAM method demonstrates conservative false discovery rate control and exhibits substantially superior performance compared to distinct when distributional changes do not involve shifts in mean (DV and DB scenarios). Conversely, in the scenarios where mean shifts contribute to the distributional change, distinct outperforms the GAM method and is even on par with conventional DA analyses like msqrob2. This observation motivated the aggregation of GAM-derived p-values with msqrob2 p-values, thereby combining the GAM methodology’s sensitivity to differences in shape with msqrob2’s established performance for detecting mean shifts.

**gamdid evaluation** Figures 6 and 7 present the FDP-TPP curves, figures 8 and 9 display the true positives at the 5% FDR threshold, comparing distinct and gamdid. The false positives are similar for both gamdid and distinct and are displayed in the supplementary materials in figures S1 and S2. gamdid’s performance is comparable to distinct for pure mean shifts (DE), demonstrates modest improvements for distributional changes that involve location shifts (DM, DP), and substantial improvements for distributional changes in shape (DV, DB).

**Figure 6:**
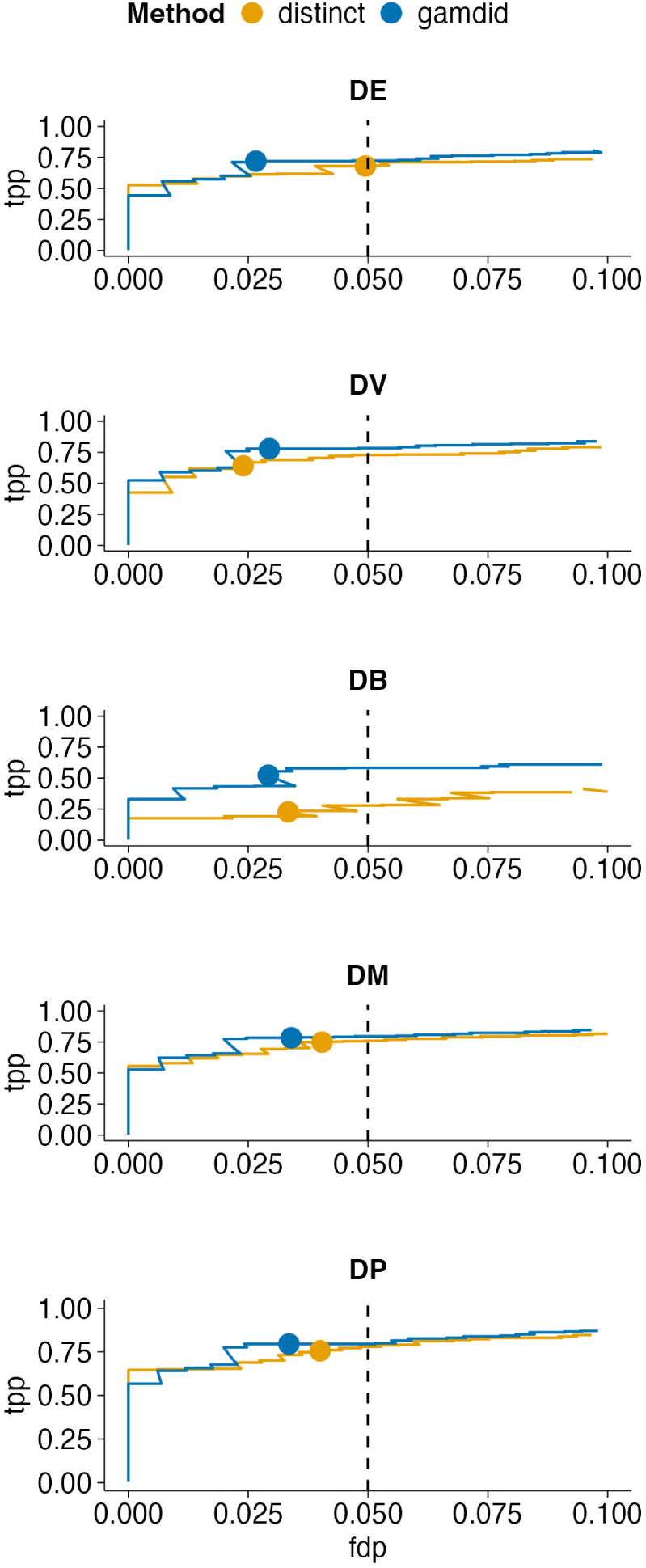
Performance of gamdid on the Leduc dataset FDP-TPP curves comparing the full gamdid method (combining GAM and msqrob2 p-values via the harmonic mean p-value) against distinct across five simulation scenarios in the Leduc dataset. By combining the local density sensitivity of the GAM with the established power of msqrob2 for mean shifts, gamdid achieves performance that is comparable to distinct for pure mean shifts (DE), shows modest improvements for distributional changes that involve a location component (DM, DP), and demonstrates substantial improvements for pure shape changes (DV, DB). gamdid maintains conservative FDR control throughout. These results demonstrate that p-value aggregation successfully bridges the complementary strengths of both components, yielding a single method competitive across the full spectrum of distributional differences.

**Figure 7:**
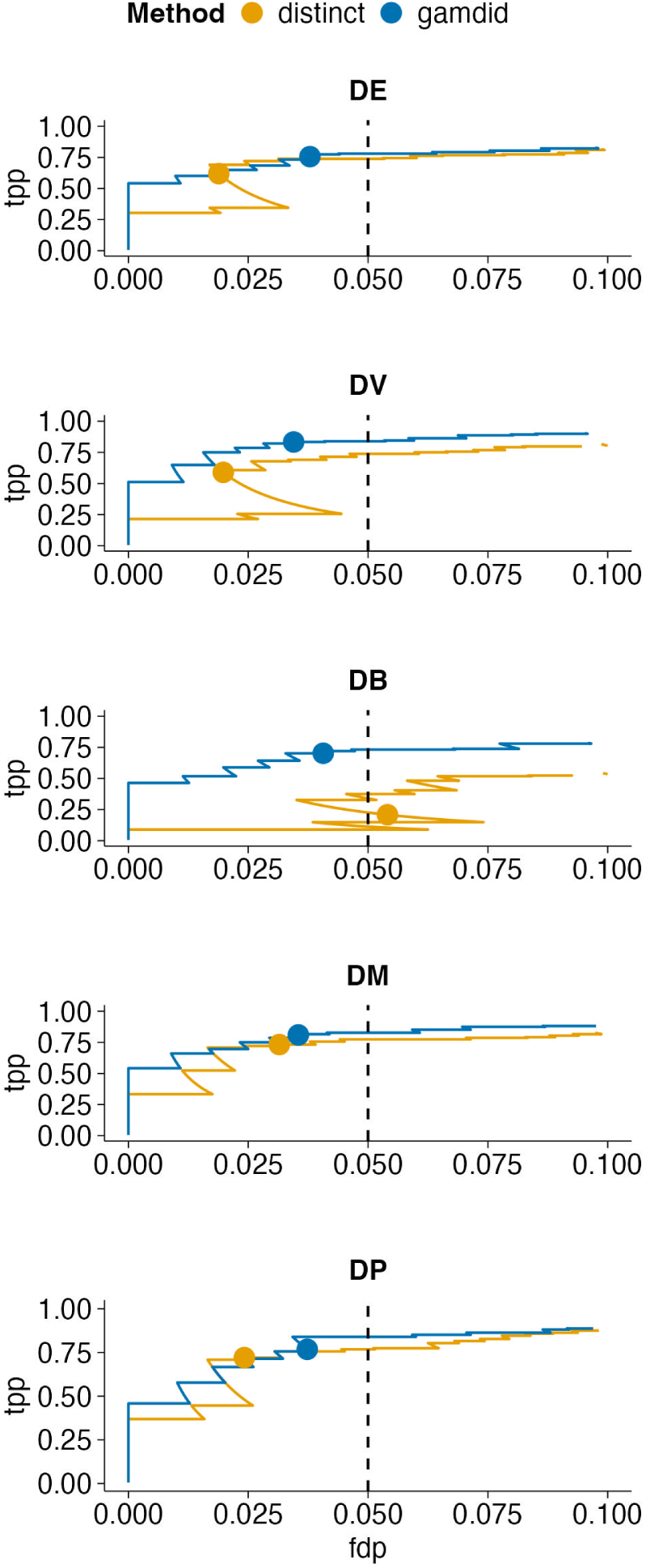
Performance of gamdid on the Petrosius dataset FDP-TPP curves comparing gamdid and distinct on the Petrosius dataset, confirming the findings from figure 6 in a second independent SCP dataset. The improvements of gamdid over distinct are more pronounced here than in the Leduc dataset. The DB scenario in particular shows a substantial improvement. These results demonstrate that gamdid’s performance advantage is robust across datasets and amplified in smaller sample size settings.

**Figure 8:**
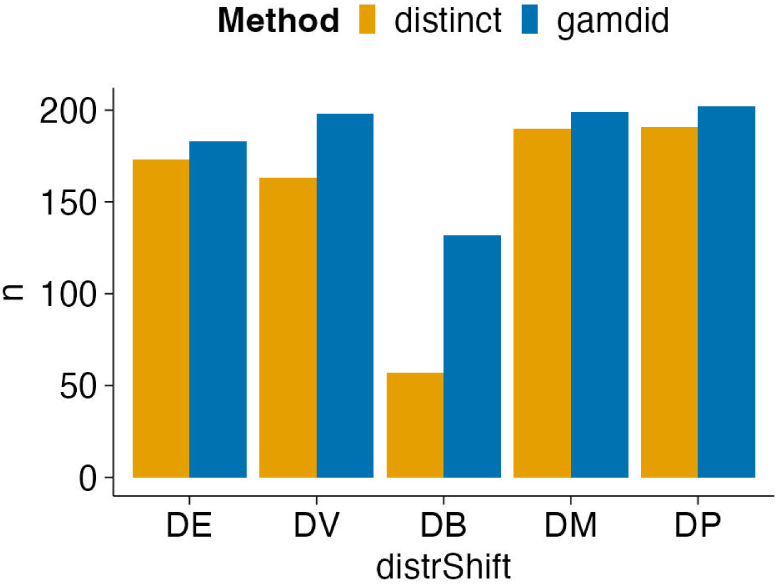
Absolute true positive counts for gamdid and distinct at 5% FDR in the Leduc dataset Bar charts displaying the number of true positives recovered at the nominal 5% FDR threshold for gamdid and distinct across the five simulation scenarios in the Leduc dataset. gamdid recovers a comparable or greater number of true positives in all scenarios, with the largest absolute gains in the DV and DB scenarios (shape-only differences). The false positive counts (figure S1) are similarly low for both methods.

**Figure 9:**
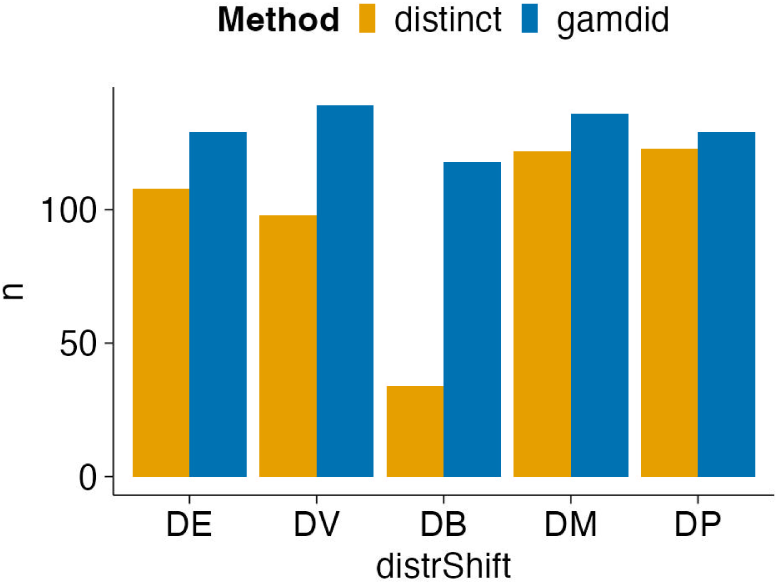
Absolute true positive counts for gamdid and distinct at 5% FDR in the Petrosius dataset Bar charts of true positive counts at the 5% FDR threshold for the Petrosius dataset, mirroring Figure 8. The advantages of gamdid are amplified compared to the Leduc results, consistent with gamdid’s higher power under smaller sample sizes. The false positive counts (figure S2) are similarly low for both methods.

Improvements are more outspoken in the Petrosius simulations, this could be attributable to the smaller sample size in the Petrosius dataset compared to Leduc. To examine this, we subsampled the Leduc dataset such that the average number of non-missing observations per group over all features matched that of the Petrosius dataset. The resulting FDP-TPP curves are provided in the supplementary materials in figure S3. Indeed, the improvements are more outspoken in the subsampled simulations compared to the Leduc simulations in figure 6. This supports the assumption that gamdid exhibits higher power under smaller sample conditions compared to distinct, which suffers from a reduced resolution of the null distribution due to a limited number of permutations.

To assess the robustness of the conclusions under less pronounced distributional differences, FDP-TPP curves for simulations with 25% reduced signal magnitudes are provided in the supplementary materials (figure S4 and S5). These results are consistent with the findings above, indicating robust performance.

### Interpretation and visualization

In contrast to competing methods, gamdid provides informative visualization of features picked up as DD. Figure 10 illustrates this functionality for a significant feature identified under the DV simulation framework. This visualization integrates GAM fits with highlighted differential regions, providing immediate visual evidence of differential variability between groups.

**Figure 10:**
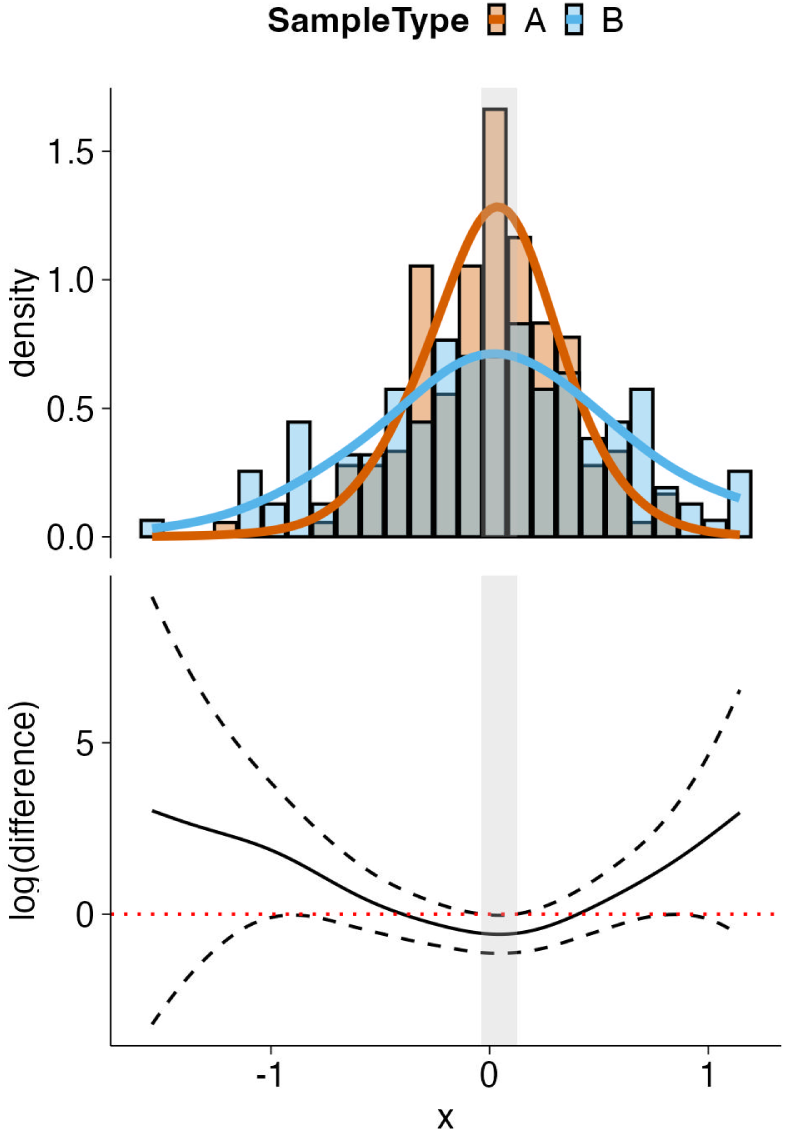
gamdid’s interpretable visualization for a DV-simulated feature in the Petrosius dataset A feature with induced differential variability (DV simulation), visualized using gamdid’s visFit function. The top panel shows the fitted GAM density curves for both groups, overlaid on the observed histograms. The bottom panel displays the difference smoother (log-ratio of densities) along with its simultaneous confidence band; the grey-shaded region identifies the specific abundance range where the distributional difference is statistically significant. Unlike competing DD methods such as distinct, gamdid does not merely return a p-value; this visualization provides an immediate visual evidence of differential variability between groups.

### Peptide level simulations

The same simulation framework was applied to peptide level assays for both the Leduc and Petrosius datasets. The resulting FDP-TPP curves are provided in the supplementary materials in figures S6 and S7 and are consistent with the protein level results.

Note, however, that the FDR control at the nominal 5% threshold is less conservative at the peptide level due to the correlation structure inherent to peptide level data, a well-known challenge in peptide level analyses. Nevertheless, this demonstrates that gamdid provides a flexible framework suitable for inferring DD at different quantification levels (precursor, peptide, proteins) in MS-based proteomics data.

**Three-group simulations** A unique advantage of gamdid is its support for omnibus testing across more than two groups, followed by post-hoc pairwise comparisons corrected for multiple testing using the stagewise testing procedure [10]. The omnibus test simultaneously assesses whether any distributional difference exists between any of the groups. No competing DD analysis method currently offers this functionality.

To evaluate this functionality, we extended the simulation framework described above to a three-group setting, randomly assigning cells into mock groups A, B, and C, with distributional signal introduced in group B under the same five simulation scenarios as above. The omnibus test was applied to this three-group comparison and the resulting FDP-TPP curves are provided in the supplementary materials in figure S8. Note that, as none of the competing DD methods supports multi-group testing, a formal benchmark comparison is not possible here.

Interestingly, the three-group setting yields higher power than the two-group comparison under equal signal strength. Note that this might arise from the default minimum observation requirement of 50 cells per group imposed by gamdid, which, in the three-group analysis, requires more observations than in the two-group analysis, leading to more powerful inference.

### Case study

We next illustrate the utility of gamdid in a case study based on the Leduc dataset.

First, we randomly divide the monocyte cells into two mock groups: X (containing 377 cells) and Y (containing 378 cells), in group Y we spike in 10% of the melanoma cells (69 cells) to simulate a subpopulation in group Y. Figure 11 shows the volcano plot of the conventional DA msqrob2 analysis twice. In the top panel, significant features of the DA analysis are marked in red, while in the bottom plot the significant features of DD methods gamdid and distinct are marked.

**Figure 11:**
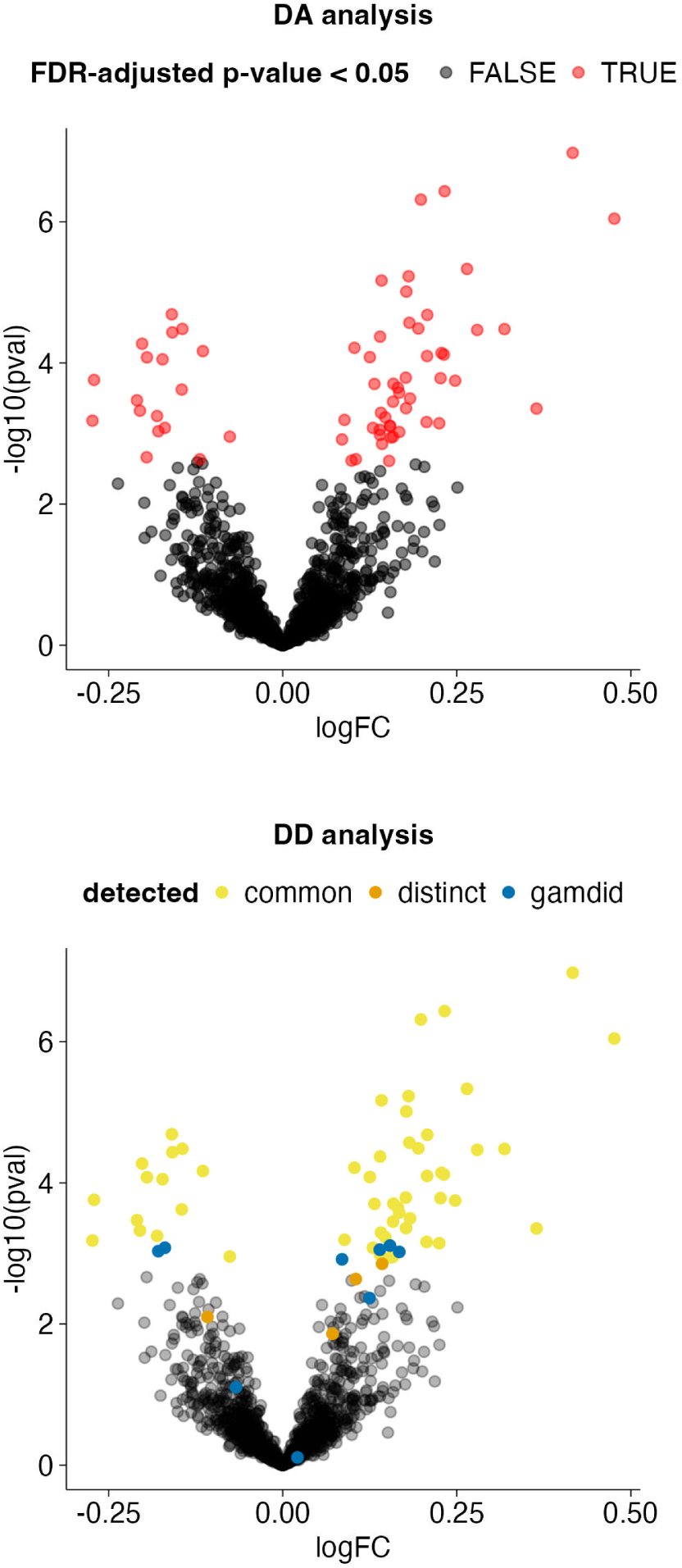
Case study: gamdid detects distributional changes missed by conventional DA analysis A controlled spike-in experiment based on the Leduc dataset: monocyte cells are split into two mock groups (X: 377 cells; Y: 378 cells), and 10% melanoma cells (69 cells) are spiked into group Y to simulate the emergence of a subpopulation. Both panels display the same msqrob2 volcano plot. Top: Features found significant by conventional DA (msqrob2) are highlighted in red. Bottom: The same volcano plot, now coloured by DD method detection: yellow = detected by both distinct and gamdid; orange = detected by distinct only; blue = detected by gamdid only. Both DD methods detect the majority of DA features, while also prioritizing some features missed by conventional DA methods.

Figure 12 presents a method comparison for the significant features in the spike-in case study. Overall, the methods produce similar results with 54 features found as significant by all three.

**Figure 12:**
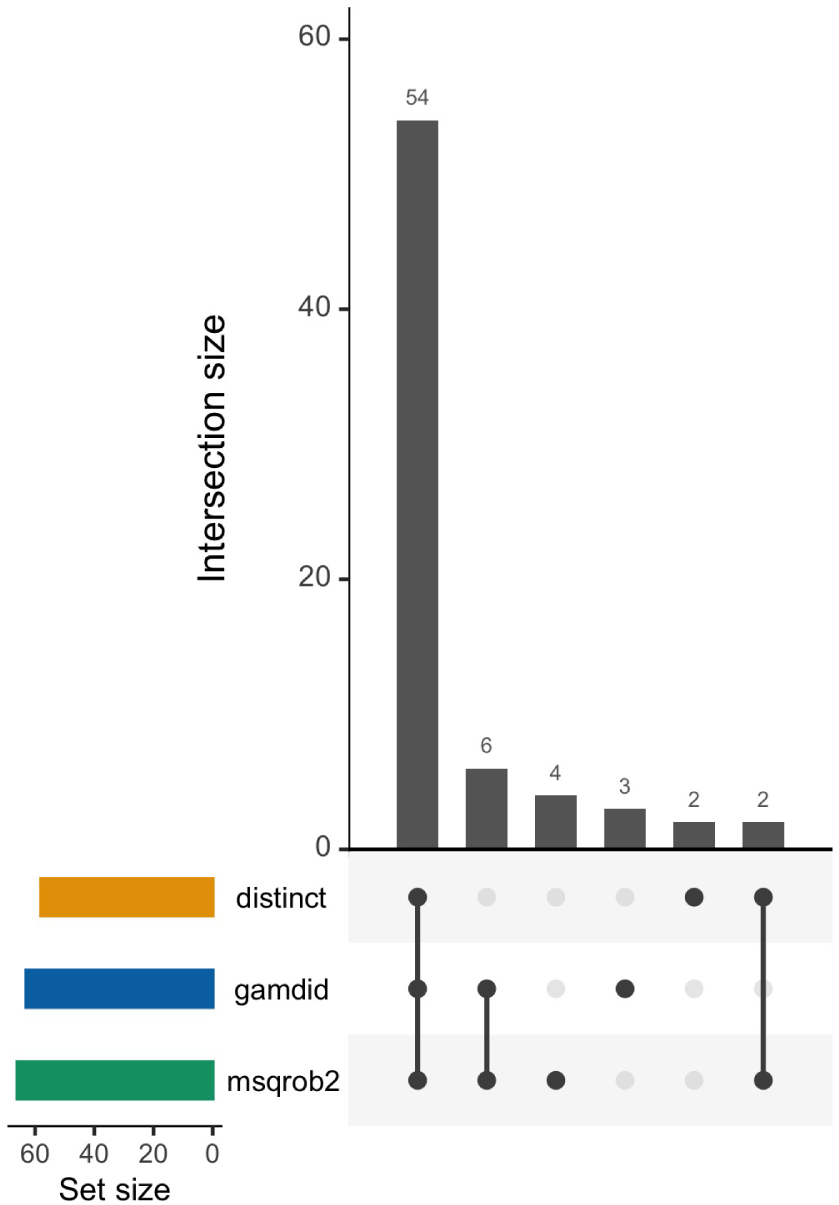
Method comparison in the spike-in case study Upset plot showing the overlap in significant features between distinct, gamdid and msqrob2 for the spike-in case study. The horizontal bars (left) show the total number of significant features per method (Set size); the vertical bars (top) show the size of each intersection, with the dot matrix below indicating which methods contribute to each intersection. Overall, the methods produce similar results with 54 features found as significant by all three. Interestingly, the 6 features picked up by the conventional DA method (msqrob2) and missed by gamdid are borderline significant for msqrob2 and borderline non-significant for gamdid (this is not the case for distinct, see table S1), which means that only gamdid largely preserves the ranking of DA features. Moreover, gamdid prioritizes 3 features that are missed by msqrob2 (see table S2).

Only six features are picked up by msqrob2 and missed by gamdid. These features are borderline significant for msqrob2 and borderline non-significant for gamdid, which is illustrated in table S1 in supplementary materials.

Furthermore, gamdid prioritizes three features that msqrob2 and distinct fail to detect (see table S2 in supplementary materials). Figure 13 shows the gamdid visualization of one of these three features, displaying the mock groups (top) and the spike-in study (bottom).

**Figure 13:**
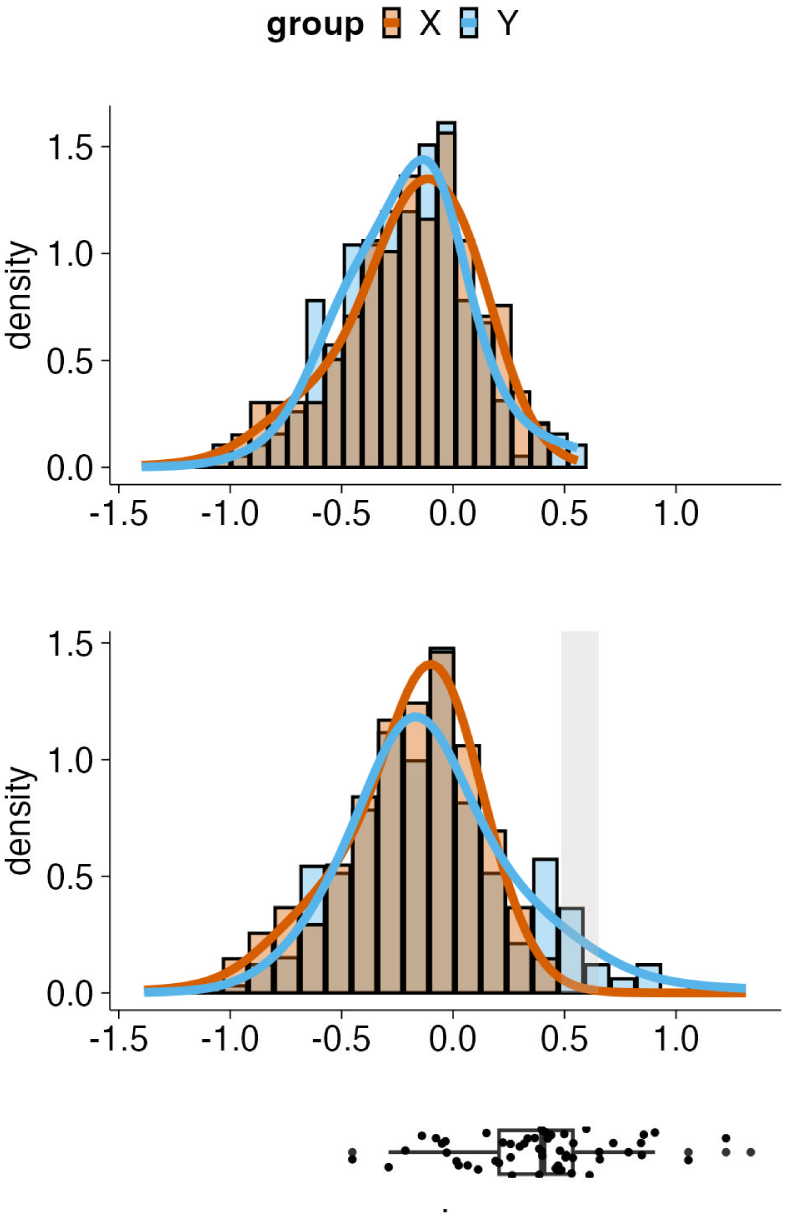
gamdid localizes the spike-in signal to the correct region of the distribution Visualization of one of the three features exclusively detected by gamdid in the spike-in case study. **Top**: The mock comparison (no spike-in), confirming that the two mock groups X and Y are distributionally equivalent before spike-in. **Bottom**: The boxplot below the figures demonstrates where the spike-in melanoma cells are located. After spiking in 10% melanoma cells into group Y, a statistically significant differential region (grey shading) appears in the right tail of the distribution, precisely where the melanoma cells were added. This result powerfully validates gamdid’s localization approach: not only does gamdid detect the distributional change, it correctly identifies its location, providing interpretable visualization that is unavailable in any competing DD method.

The boxplot below the gamdid visualizations indicates where the spiked-in melanoma cells are located within the distribution of group Y. The spiked-in cells align perfectly with the detected differential region. This clearly illustrates the advantage of gamdid over other DA and DD methods: its functionality to localize changes in distribution and combine it with an interpretable visualization.

The gamdid visualization of the two features only detected by msqrob2 and distinct, respectively, are provided in supplementary materials in figures S9, S10, S11, and S12. Interestingly, features detected by all three methods also illustrate the utility of gamdid. Four representative features are provided in the supplementary materials in figure S13, which clearly show that these features are DD due to the subpopulation of spike-in cells rather than DA.

## Discussion

### Heterogeneity in single cell data

Because of gamdid’s flexible approach, it captures the heterogeneity of single cell data that are by design missed by conventional DA methods. In the case study, gamdid identifies three features that are not detected by conventional DA methods as the distributional differences do not involve a sufficiently large mean shift.

More importantly, even the 54 features detected by both DA and DD methods further emphasize the need to account for heterogeneity in single cell data. Indeed, for the majority of these features, the distributional difference extends beyond a pure mean shift. This is illustrated by gamdid’s visualization which reveals the important nuances of these more complex distributional patterns.

Four representative examples are provided in the supplementary materials in figure S13. Notably, in all four examples the detected differential regions align closely with the regions of the distribution where melanoma cells were spiked in, underscoring the utility of gamdid’s visualization.

### Ranking of significant features

Notably, the ranking of significant features is of increasing interest in proteomics, and gamdid offers a key advantage in this regard. For DA features, the gamdid ranking is very similar to top-performing DA methods (which is not the case for distinct, see table S1). Moreover, gamdid can also prioritize interesting features that are completely overlooked by DA methods. This is illustrated in table S2, which shows that two out of the three features exclusively detected by gamdid are assigned high p values by msqrob2.

**Conceptual difference between** gamdid **and** distinct The simulation results reflect a fundamental difference in how distinct and the GAM methodology characterize distributional differences between groups. distinct operates on empirical cumulative distribution functions (ECDFs) and quantifies the mean difference between the ECDFs of two groups. By design, this mean ECDF difference is sensitive to sustained, global displacements of probability mass. The GAM-based approach, however, models the density function of each group. Because density estimation is inherently local, the GAM is particularly well-suited to detect distributional changes that are concentrated in specific regions of the distribution.

Figure 14 illustrates this using a feature from the Petrosius DB simulations that exhibits a local change without substantially displacing the bulk of the probability mass. In this case, the mean difference between the ECDFs of the two groups remains modest, resulting in reduced power for distinct, whereas the GAM-based method is able to successfully identify the differential region.

**Figure 14:**
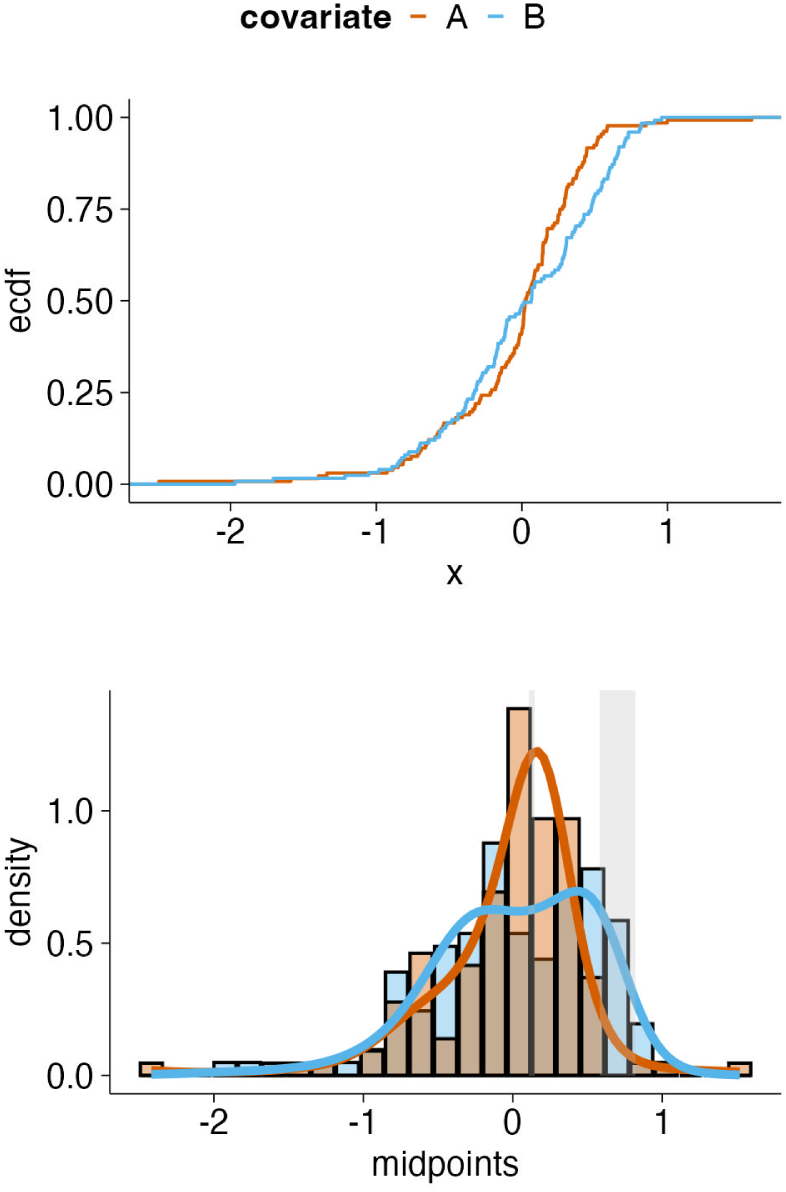
Why gamdid outperforms distinct for shape differences Comparison of how distinct and gamdid characterize a feature with differential component means (DB) from the Petrosius simulations. The feature was detected by gamdid (FDR adjusted p-value = 0.012) and not by distinct (FDR adjusted p-value = 0.278). **Top**: The ECDF curves used by distinct. The mean ECDF difference between groups is modest because the induced bimodality does not produce a sustained, global displacement of probability mass. **Bottom**: The gamdid visualization reveals the local bimodality clearly, and the difference smoother identifies the specific regions where the densities diverge. This contrast illustrates the fundamental conceptual difference between the two methods: distinct is more sensitive to sustained, global ECDF displacements (as occur with location shifts), while the GAM’s density-based, inherently local approach is better suited to detecting concentrated, shape-based differences. This complementarity is precisely what motivates gamdid’s p-value aggregation strategy.

Conversely, distributional changes like mean shifts produce large, sustained differences between the ECDF’s of two groups and are therefore readily detected by distinct. The GAM-based method, however, exhibits reduced power in such settings, as mean shifts constitute diffuse, global changes that do not concentrate in any particular local region of the distribution. This behavior is illustrated in figure 15, which presents a feature from the Petrosius DE simulations. This example shows an accumulated difference between the ECDFs, while the GAM method is not able to detect differential regions.

**Figure 15:**
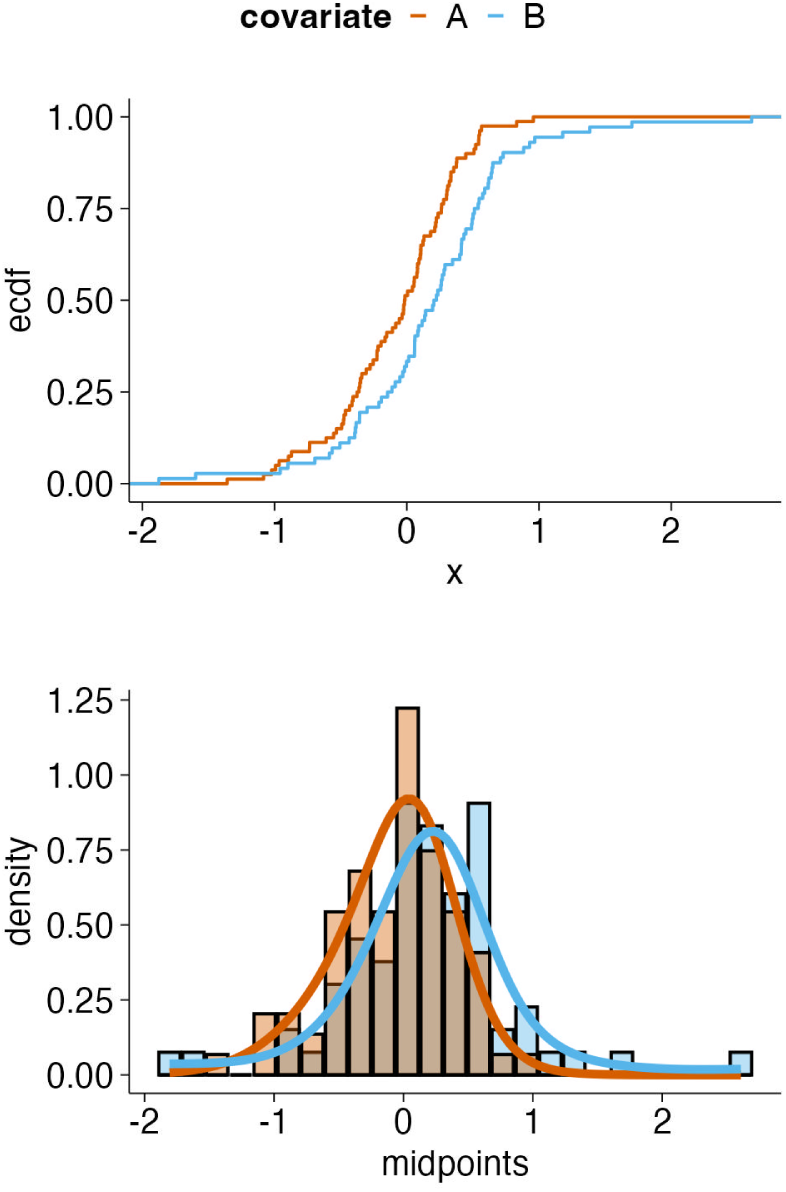
Why gamdid aggregates with msqrob2 for mean shifts Comparison of how distinct and gamdid characterize a feature with a pure mean shift (DE) from the Petrosius simulations.The feature was detected by distinct (FDR adjusted p-value = 0.020) and not by the standalone GAM method before aggregation (FDR adjusted p-value = 0.227). **Top**: The ECDF curves show a large, sustained horizontal displacement between groups. **Bottom**: The gamdid density visualization shows overlapping GAM fits with no identifiable differential region (no grey shading), reflecting the GAM’s reduced power for diffuse, global distributional changes that are not concentrated in any specific region. This example again motivates the aggregation of GAM p-values with msqrob2 p-values: by combining the local sensitivity of the GAM for shape changes with the sensitivity of msqrob2 for location shifts, gamdid achieves good performance across all types of distributional differences. In this example the gamdid FDR adjusted p-value is 0.029.

This motivates the aggregation of GAM-derived p-values with those from msqrob2, which is specifically designed to detect mean shifts with high sensitivity.

**P-value aggregation** The aggregated p-value in gamdid combines the local sensitivity of the GAM with the established power of msqrob2 for mean-based inference, ensuring that gamdid is well-suited to detect any form of distributional differences. As expected, a modest loss of power relative to each individual component is observed in scenarios where only one of the two components contributes meaningfully: the DE scenario for the GAM component, and the DV and DB scenarios for msqrob2. This is an inherent consequence of p-value aggregation when one of the contributing p-values carries no signal. In scenarios where the distributional change involves both a mean shift and a difference in shape, however, gamdid combines the strength of both methods and outperforms each method separately, demonstrating the practical benefit of the aggregation strategy.

**FDR control** gamdid achieves conservative FDR control at the nominal 5% threshold throughout all simulations.

### Computational efficiency

To evaluate computational efficiency, the runtime of both gamdid and distinct was measured on the spike-in case study dataset, comprising 1316 features and 824 cells. Each method was executed on a single core, and runtimes were averaged across ten independent runs. The mean runtime for gamdid was 16 seconds, whereas distinct required 256 seconds on average, indicating that gamdid achieved a speedup of more than fifteen-fold (see figure 16).

**Figure 16:**
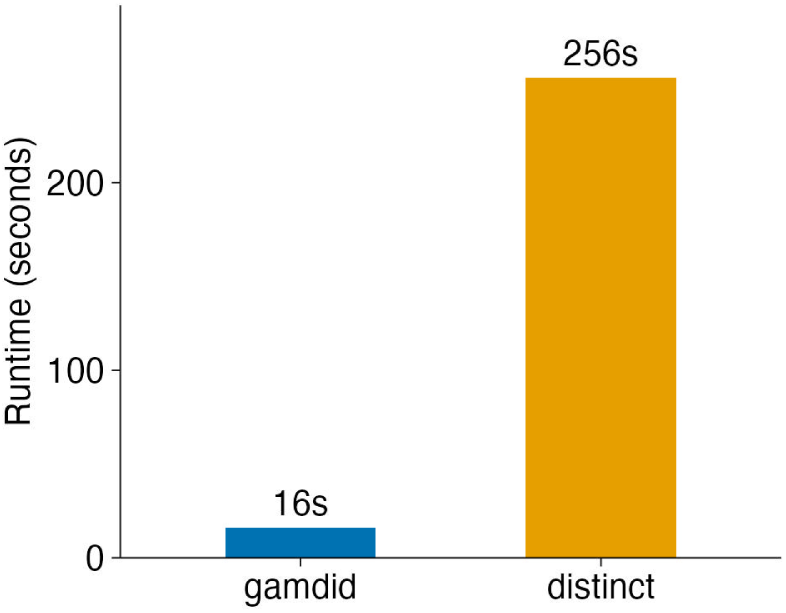
Computational efficiency of gamdid and distinct. Mean runtime for gamdid and distinct on the spike-in case study dataset (1316 features, 824 cells), each run on a single core and averaged across ten independent runs. gamdid completed in 16 seconds on average, whereas distinct required 256 seconds, corresponding to a more than fifteen-fold speedup.

This can be attributed to two key factors. First, gamdid does not rely on permutation-based inference as distinct does. Second, as distinct cannot handle missing values, all missing observations must be removed on a per-feature basis prior to analysis.

### Outlook

The development of gamdid represents a significant advance toward a more comprehensive interpretation of the information in single-cell proteomics data. By extending differential analysis beyond mean-based comparisons, gamdid enables the systematic investigation of distributional changes and thereby facilitates a more detailed characterization of cellular heterogeneity. This functionality uncovers biologically relevant variation that has remained largely unexplored by existing data analysis frameworks.

As single-cell proteomics technologies continue to mature and datasets grow in size and complexity, the ability to detect and interpret subtle distributional differences will become increasingly important. We therefore anticipate that gamdid can find applications not only in the comparison of cell types and conditions, but also in the identification of subpopulations, or the characterization of transitional cell states.

More broadly, the framework presented here is not intrinsically limited to proteomics and could, in principle, be adapted to other single-cell modalities, including single-cell transcriptomics, metabolomics and flow cytometry, where distributional heterogeneity likewise constitutes an important source of biological information. As such, gamdid has the potential to support a broader methodological shift toward distributional approaches in single-cell data analysis, enabling a more comprehensive characterization of cellular variation beyond changes in average abundance levels.

## Methods

We first detail the density estimation approach and the Poisson generalized additive model used. We subsequently describe the inference procedure and the p-value aggregation strategy. Next we elaborate on the post-hoc pairwise comparisons and stagewise testing procedure, followed by the approach for identifying regions of differential density. The section closes with a description of the covariate adjustment procedure and the two datasets used throughout the manuscript.

### Model and inference

The statistical methodology for comparing distributions is developed for a single feature (in this case proteins or peptides). The method is then simply repeated over multiple features and the corresponding p-values are corrected for multiple testing using conventional Benjamini-Hochberg false discovery rate (FDR) method [11] or stagewise testing [10].

### Density estimation through Poisson regression

Following the approach of Lindsay [5], we reformulate density estimation in terms of Poisson regression by discretizing the sample space. We partition the sample space *Y* into K disjoint bins *Yk* of equal binwidth *bw*.

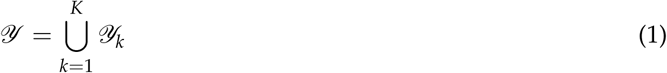

The data ***y*** = (*y*_1_, *y*_2_, …*y_n_*)*^T^* are summarized into bin counts ***s*** = (*s*_1_, *s*_2_, …*s_K_*)*^T^* where each count *s_k_* represents the number of observations falling into the *k*-th bin:

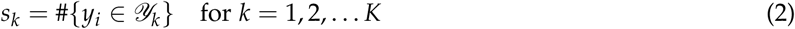

The sum of these counts equals the total number of observations *n*. Conceptually, this process is equivalent to constructing a histogram of the data, where each bin Y*_k_*is associated with a count *s_k_* and represented by a midpoint *mid_k_*.

We consider *s_k_* to be independent Poisson observations with expected values *µk*

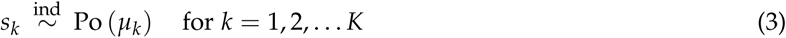

Consequently, the problem of density estimation is recast in spline-based Poisson regression of the bin counts *s_k_* in function of the bin midpoints *mid_k_* [12].

### Poisson generalized additive model

The expected bin counts *µk* are modelled using Poisson regression implemented in the existing R package mgcv [13] for generalized additive models (GAM).

The linear predictor of a standard GAM model with one smoother ( *f* (*mid_k_*)) is given by equation 4.

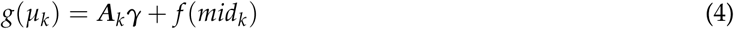

with ***A****_k_* the k-th row of the parametric *K* × *G* model matrix ***A*** and ***γ*** the parametric part of the model parameter vector. For Poisson distributed data, the link function *g* is a log function.

The relationship between *log*(*µk* ) and *mid_k_*, *f* (*mid_k_*), is modeled using penalized thin plate regression splines (see equation 5) that provide a basis function expansion, i.e.

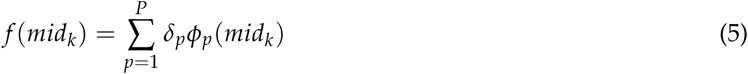

where *ϕ_p_*(.) are thin plate spline basis functions, *δ_p_* are the corresponding smooth basis parameters to be estimated. Let ***β*** = (***γ***, ***δ***) where ***δ*** = (*δ*_1_, . . . , *δ_P_*)*^T^*.

Let **Φ** be the *K* × *P* model matrix of the basis function evaluated in the midpoint: *ϕ_p_*(*mid_k_*). We then combine ***A*** and **Φ** in the *K* × (*G* + *P*) model matrix ***X*** = (***A*** : **Φ**)

***β*** is estimated by maximizing the penalized log-likelihood (see equation 6)

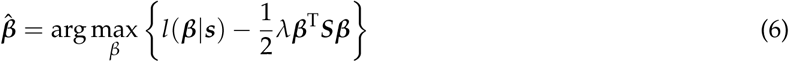

The optimization is typically performed using penalized iteratively reweighted least squares (PIRLS). Here, *l*(***β***|***s***) is the log likelihood of the data given ***β***, *λ* is the smoothing parameter that balances goodness-of-fit with smoothness, and ***S*** is the penalty matrix quantifying the wiggliness of the basis functions that has zeros on the rows and columns involving parameters for ***A*** [14]. We exploit the link between Bayesian statistics, smooth functions, and mixed models to estimate this smoothing parameter *λ* using the REML method [15], [16].

To approximate the effective degrees of freedom (edf) of the model, we define the influence matrix ***H*** in equation 7.

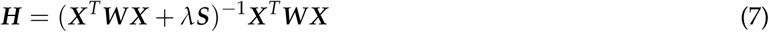

with ***W*** the diagonal weight matrix from the final PIRLS iteration, where *W_kk_* = *µk* = *exp*(***X*** *^T^ **β***^ ) and ***X****_k_* is the k-th row of ***X***.

In the GAM literature 2***H*** − ***HH*** is the bias corrected influence matrix (the bias arises due to smoothing), so its trace can be seen as the effective degrees of freedom of the bias corrected model (see equation 8) [17].

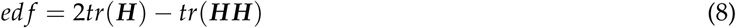

### Comparing distributions across groups

We build on this framework to detect distributional differences across multiple groups. Let *c* be the groups of grouping variable *C*.

Next, the overall sample space *Y* is partitioned for all groups *c* into K disjoint bins *Yk* of equal width *bw* (as in equation 1), ensuring identical bins (and thus midpoints *mid_k_*) for all groups *c* ∈ *C*, as illustrated in figure 2 panel A. For each group *c*, a count vector ***s_c_*** = (*s_c_*1 , *s_c_*2 , …, *s_c_K* ) is obtained, where *s_c_k* is the number of observations from group *c* in the k-th bin.

The full model allows for different distributions across groups, with the linear predictor specified in equation 9.

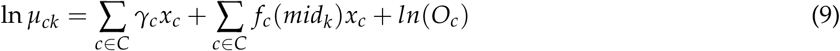

where each group *c* has its own intercept *γ_c_* and smooth function *f_c_*, with identical smoothing parameter *λ* among all groups.

The offset *O_c_* is defined as

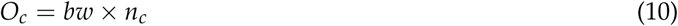

where *n_c_* is the total number of observations in group *c*. This allows for normalization of different sample sizes across groups and ensures that the fitted smooth functions *exp*(*γ_c_* + *f_c_*(*mid_k_*)) can be directly interpreted as probability density.

Conversely, the null model assumes a common density function across all groups of *C*. The linear predictor is modeled as in equation 11.

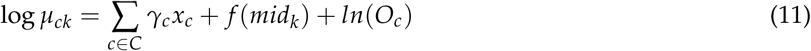

where *x_c_* is a dummy variable for the groups of covariate *C*, *µck* is the expected count for group *c* in bin k, and the smooth function *f* (mid*_k_*) is shared across all groups. *O_c_* is an offset term.

### Inference for distributional differences

Two approaches were considered for testing whether distributions differ across groups.

• Generalized likelihood ratio test (GLRT)

This approach is possible by comparing the full and the null model. However, it requires fitting two models and does not directly provide pairwise comparisons for a multi group setting. Therefore, we provide a Wald-type test below and refer the reader to the supplementary materials for the GLRT test.

• Wald-type test

A more practical approach involves Wald-type tests on the parameters of the smooth functions of the full model (see equation 9). This method builds on tests proposed by Simon Wood, which assess whether a smoother significantly deviates from zero, *H*_0_ : *f_c_* = 0 [18].

We extend this idea to test whether multiple smooth functions are identical across *G* groups. Specifically, we test the null hypothesis *H*_0_ : *f_c_*1 = *f_c_*2 = · · · = *f_c_G* . This hypothesis can be reformulated as a Wald test on the corresponding basis function parameters. Indeed, equality of smooth functions *f_c_*1 = *f_c_*2 = · · · = *f_c_G* implies equality of their associated parameter vectors ***β****c*1 = ***β****c*2 = · · · = ***β****cG* if basis functions are shared across all groups.

To test this hypothesis, we define contrasts on the full vector of basis function parameters ***β***^ full = (***β***^ *c*1 , . . . , ***β***^ *cG* ) from the full model in equation 9. The null hypothesis can then be expressed as *H*_0_ : ***Lβ***full = 0 where ***L*** is a contrast matrix encoding *G* − 1 independent differences between parameter vectors, i.e. (***β****c*1 − ***β****cg* = 0 for *g* = 2, · · · , *G*).

The corresponding Wald statistic is then given by equation 12.

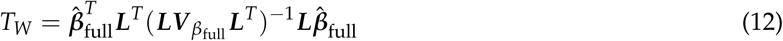

where *V_βfull_* = ((**X**^T^**WX** + *λ***S**)^−1^ is the covariance matrix for ***β*** . *T_W_* is asymptotically *χ*^2^ distributed.

The degrees of freedom *z* correspond to the total effective degrees of freedom (edf) associated with the *G* − 1 smooth functions that are dropped under *H*_0_. As a conservative estimate, we define *z* as the sum of the *G* − 1 largest edf values.

The p-value obtained from the *χ*^2^ distribution of *T_W_* reflects any form of distributional deviation between groups: mean shifts, differences in variance, modality, skewness… In what follows, we refer to this as the GAM p-value.

### P-value aggregation

While the GAM p-value is sensitive for a wide range of distributional differences, it exhibits limited power in detecting pure mean shifts. To improve sensitivity to such shifts, we complement the GAM p-value with a p-value derived from the msqrob2 framework, which is specifically designed to detect mean shifts in proteomics data using a robust linear regression through a mixed model framework [6].

The GAM p-value and the msqrob2 p-value are aggregated using the harmonic mean p (HMP) method. The HMP method is particularly suited for combining correlated p-values [7]. This is inherently the case since mean shifts are one aspect of overall distributional changes. The resulting aggregated p-value, termed the ’feature-level aggregated p-value’, provides a single, comprehensive measure of evidence against the null hypothesis of no distributional difference among groups for that feature.

### Post-hoc pairwise comparisons

If covariate *C* comprises more than two groups, post-hoc pairwise comparisons can be conducted. For each of the C(C − 1)/2 pairs of groups (where C is the number of groups in *C*), a pairwise GAM p-value is calculated using the Wald test framework (see equation 12) with an appropriate contrast matrix ***L***. Similarly, pairwise msqrob2 p-values are computed. These pairwise p-values are then aggregated for each group-to-group comparison using the HMP method, yielding ’pairwise aggregated p-values’, each representing the evidence against the null hypothesis of identical distributions for that specific pair of groups.

### Stagewise testing

A stage-wise approach is employed to effectively manage the multiple hypotheses arising from feature-level and subsequent pairwise comparisons [10].

- Stage 1 (feature level FDR control): The feature level aggregated p-values are adjusted across all features using the Benjamini-Hochberg (BH) procedure to control the family wise error rate (FDR) at 5%.
- Stage 2 (Pairwise FWER control) is only conducted if stage 1 results in a significant p-value. The pairwise aggregated p-values are adjusted for multiple testing across the C(C − 1)/2 pairwise hypotheses within a feature to control the family wise error rate (FWER) on the BH-adjusted significance level (i.e., the FDR q-value) of the first stage [19].

### Identifying regions of differential density

For pairs of groups exhibiting significant distributional differences, the specific regions contributing to these differences can be pinpointed. This is achieved by constructing a difference smoother *d*^, representing the estimated difference in log-densities between the groups, and its associated simultaneous confidence bands. Consider the comparison between two groups: *c*_1_ and *c*_2_ from the groups of *C*. The difference smoother *d*^ is evaluated across a grid of *N* points: *x_n_* for *n* = (1, …, *N*) spanning the range of *mid_k_* values (see equation 13).

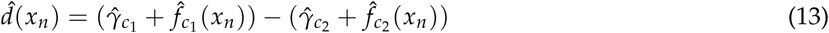

Note that, *d*^(*x_n_*) can also be expressed as a linear combination of the full parameter vector ***β***^ full. Let ***x****_di f f_* be the contrast row vector for the comparison of *c*_1_ and *c*_2_ so that the difference smoother can be defined as in equation 14.

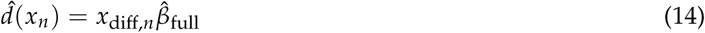

The corresponding pointwise standard error is subsequently computed in equation 15.

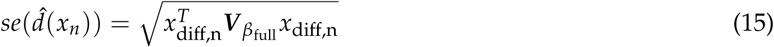

where ***V*** *_β_*full is the covariance matrix for *β*^full.

A 1 − *α* pointwise confidence interval for *d*^(*x_n_*) is then given by *d*^(*x_n_*) ± *z_α_*_/2_ ∗ *se*(*d*^(*x_n_*)) where *z_α_*_/2_ is the 1 − *α*/2 quantile of the standard normal distribution.

To enable valid inference over the entire difference smoother *d*^ simultaneously, the pointwise confidence intervals need to be adjusted for multiple testing. Essentially, this is equivalent to adjusting for the multiple pointwise tests conducted along the grid. As a computationally efficient approach, we use the Šidák multiple testing correction to transform the pointwise intervals into simultaneous confidence bands [20]. The Šidák correction will be conservative because the points are positively correlated [21].

Regions where the simultaneous band does not encompass zero, can then be used to prioritize DD regions. The 95% simultaneous confidence interval for *d*^(*x_n_*) then becomes *d*^(*x_n_*) ± *z*∗ *se*(*d*^(*x_n_*)), where the adjusted critical value *z*∗ is *z*1 −(1−*α*overall) 1/*M* . *M* denotes the effective number of independent locations along the grid, which is conservatively estimated as the maximum of the effective degrees of freedom associated with the two smooth functions being compared *f*^*c*1 and *f*^*c*2 . This choice is motivated by the equivalence between such grid tests and omnibus Wald-type tests on the basis function parameters (see equation 12).

Similar to the previous section, a stagewise testing approach is used to correct for multiple testing along the features. This means that *α*overall corresponds to the BH adjusted significance level of the first stage to control the feature-level FDR at 5% [10].

Note, however, that we do not provide simultaneous confidence intervals that account for multiple testing along features as well as within feature to address multiple comparisons between groups. Indeed, this would require the extension of the stagewise testing procedure towards multiple stages.

### Covariate adjustment

In addition to the covariate of interest, gamdid allows users to account for confounding variables through a nuisance formula, enabling applications such as batch correction in multi-batch experimental designs. Both the covariate of interest and the nuisance formula are combined into one model in the msqrob2 analysis, and an appropriate contrast is constructed to test for distributional differences in the covariate of interest.

To remove the influence of nuisance covariates from the GAM-based inference, partial residuals with respect to the covariate of interest are computed from the msqrob2 fit and subsequently used as input to the GAM analysis. This two-step approach is required due to the binning strategy: as a single bin may contain observations that correspond to different groups of the nuisance covariate, direct covariate adjustment within the GAM framework is not feasible.

Note, however, that the uncertainty associated to the regression of the nuisance covariates in step one are not accounted for, so the resulting p-value should be interpreted with caution.

### Data

- leduc2022_pSCoPE: This data was acquired using pSCoPE technology: a TMT-18plex protocol with prioritized data-dependent acquisition (DDA) carried out by a Thermo Scientific Q-Exactive mass spectrometer [1]. The MS data were quantified and identified using MaxQuant version 1.6.17 [22]. We performed quality control by filtering low quality features and filtering cells with a median coefficient of variation greater than 0.6, calculated using the *medianCVperCell* function of the scp package [8]. The PSM intensities were log2-transformed for variance-stabilization, median centered and summarized to protein-level using median summarization per MS run. Subsequently, samples were normalized through median centering and proteins with more than 95% missingness were filtered out, resulting in a dataset with 776 melanoma cells and 755 monocytes profiled across 2120 proteins.
- petrosius2023_mES: This data originates from a label-free scp experiment conducted on mouse embryonic stem cells cultured under two distinct conditions: a serum-free 2i condition (m2i) containing cytokine LIF and inhibitors targeting the MEK and GSK3 pathways, and a serum condition (m15) only containing cytokine LIF [9]. Data acquisition was performed using an Orbitrap Eclipse Tribrid mass spectrometer operating in data-independent (DIA) acquisition mode. The resulting mass spectrometry data were subsequently analyzed by the authors using Spectronaut v17 [23]. We performed quality control by removing cells with fewer than 500 detected peptides or a median intensity lower than 11. Features with more than 95% missing values were also removed. The detected intensities were log2-transformed for variance-stabilization, and the intensities of each cell were centered with the median log2-intensity of the corresponding cell. Subsequently, peptides were summarized to protein-level through median polish summarization. The resulting dataset contains 295 m15 and 269 m2i cells with 2492 proteins.

Preprocessed protein-level summarized assays from both datasets served as the foundation for our simulations. From the Petrosius dataset, we selected the 269 m2i-condition cells, while from Leduc, we selected the 755 monocytes (in both cases following removal of non-informative cells during preprocessing). Within each selected subset, no systematic biological differences are expected between cells. Consequently, random assignment of cells into two mock groups (A and B) yields a dataset suitable as a mock basis for a controlled two-group comparative analysis.

### Simulation framework

The benchmark simulations are based on protein-level mock data from both the Leduc and Petrosius datasets, as described in the Results section. For each dataset, mock groups A and B were formed by random assignment of cells, and distributional signal was introduced in group B for 20% of randomly selected features. Five types of distributional signal were considered, following the framework of Korthauer et al. (2016) [3]:

- **DE** A shift equal to 40% of the feature’s standard deviation is added to all observations in group B.
- **DV** Independent Gaussian noise with mean zero and standard deviation equal to 120% of the feature’s standard deviation is added to all observations in group B, thereby increasing the spread of the distribution without altering its mean.
- **DB** For 50% of the observations in group B, a value of 90% of the feature’s standard deviation is added; for the remaining 50%, the same value is subtracted. This induces a bimodal distribution in group B.
- **DM** A value of 90% of the feature’s standard deviation is added to 50% of the observations in group B, creating an asymmetric shift concentrated in one tail.
- **DP** A value of 90% of the feature’s standard deviation is added to 40% of the observations in group A and to 70% of the observations in group B, thereby inducing a difference in the relative weight of a subpopulation.

### Implementation

The methods for assessing differential distributions are implemented in the gamdid R package that is publicly available through github (github.com/statOmics/gamdid) and will be submitted to Bioconductor.

gamdid builds upon the CRAN packages mgcv [13] and harmonicmeanp [24] for fitting generalized additive models and p-value aggregation; and on the R/Bioconductor packages stageR [19] for stagewise testing and msqrob2 for detecting shifts in mean abundance [6]. Particularly, mgcv version 1.9.4, harmonicmeanp version 3.0.1, stageR version 1.32.0 and msqrob2 version 1.18.0 were used in R version 4.5.3.

## Data Availability

All data used in this manuscript are publicly available. The Leduc [1] and Petrosius [9] datasets were downloaded using the scpdata [8], [25] package from Bioconductor.

## Code Availability

All code used to prepare and analyze the datasets and produce the figures of this manuscript are available through github/statOmics/gamdid. Finally, gamdid is published as an open source R-package available through github (github.com/statOmics/gamdid) and will be submitted to Bioconductor.

## Supplementary materials

**Figure S1:**
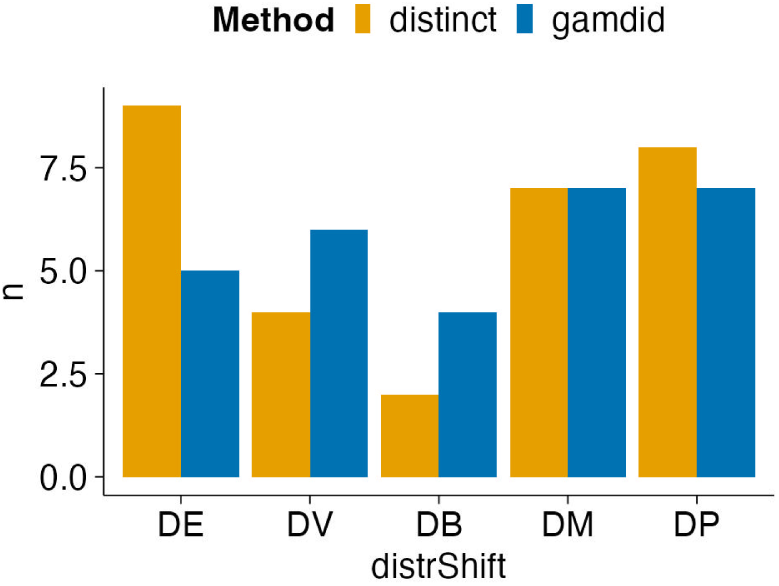
False positive counts for gamdid and distinct at 5% FDR for the Leduc dataset Bar charts displaying the number of false positives at the nominal 5% FDR threshold for gamdid and distinct across the five simulation scenarios in the Leduc dataset. Both methods maintain similarly low false positive counts across all scenarios, confirming that the gains in true positive recovery demonstrated for gamdid in figure 8 are not accompanied by inflation of false discoveries.

**Figure S2:**
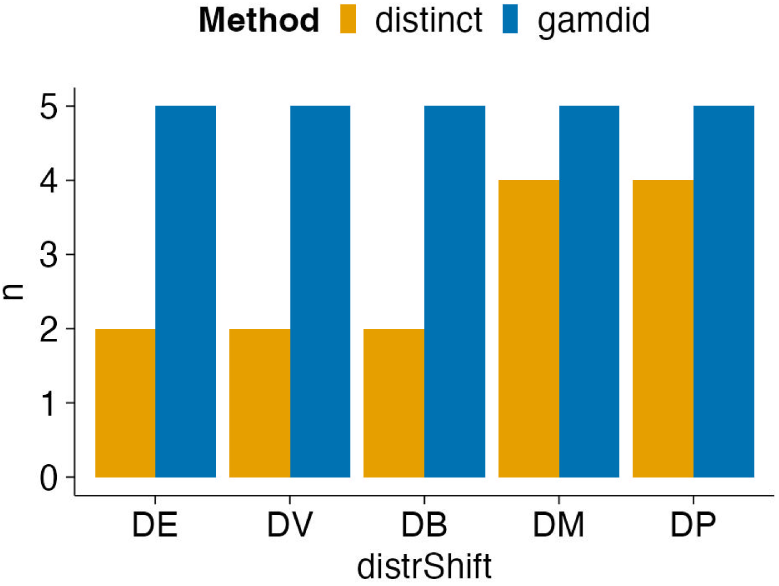
False positive counts for gamdid and distinct at 5% FDR for the Petrosius dataset Bar charts of false positive counts at the 5% FDR threshold for the Petrosius dataset. As in the Leduc dataset, false positive counts are similarly low for both methods across all five simulation scenarios, confirming that the gains in true positive recovery demonstrated for gamdid in figure 9 are not accompanied by inflation of false discoveries.

**Figure S3:**
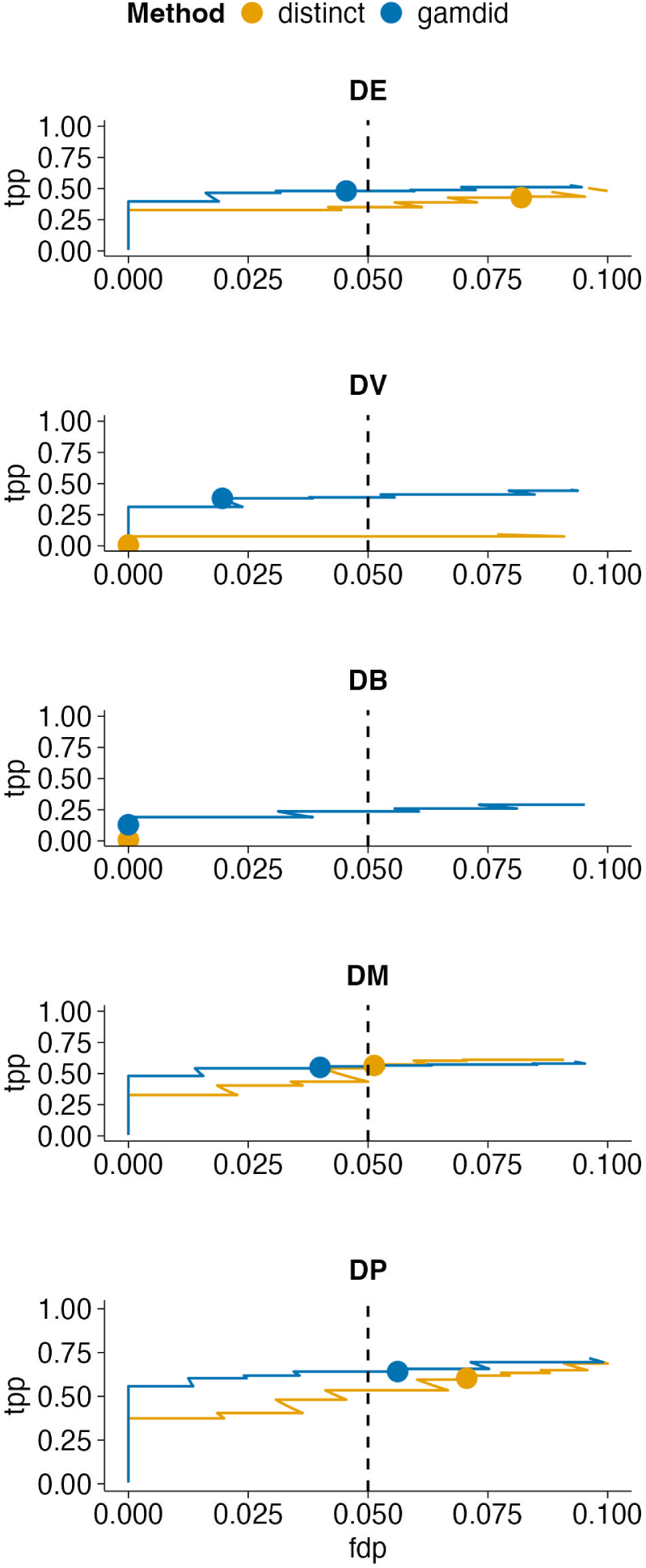
Performance of gamdid and distinct on a subsampled Leduc dataset FDP-TPP curves comparing gamdid and distinct on a subsampled version of the Leduc dataset, in which cells were subsampled such that the average number of non-missing observations per group matched that of the Petrosius dataset. This analysis directly tests whether the more pronounced improvements of gamdid over distinct observed in the Petrosius dataset (see figure 7) are attributable to the smaller sample size. The results support this hypothesis: under reduced sample size, the performance gap between gamdid and distinct increases. distinct likely suffers from reduced null distribution resolution when fewer observations are available.

**Figure S4:**
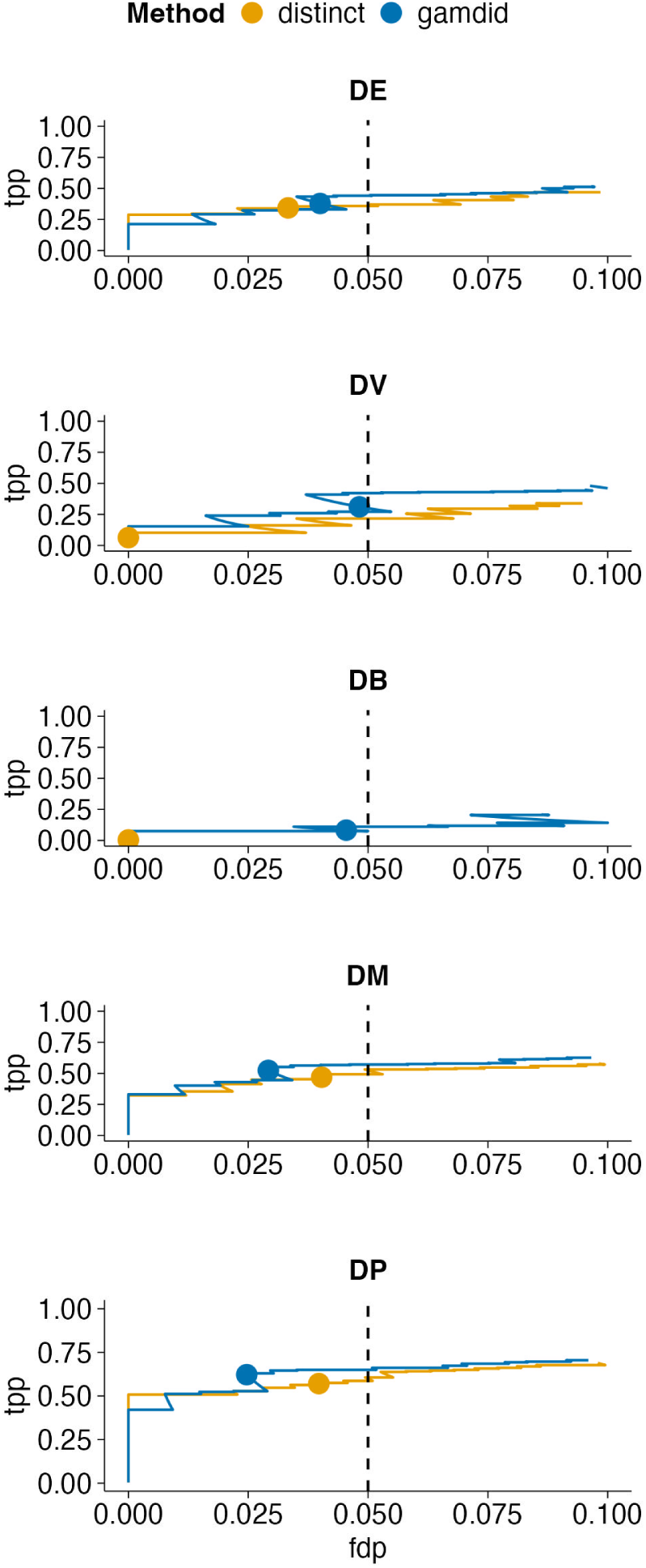
Robustness of gamdid performance under reduced signal magnitude for the Leduc dataset FDP-TPP curves comparing gamdid and distinct in the Leduc dataset under simulation scenarios with 25% reduced signal magnitudes (DE: a shift of 30% instead of 40% of the feature’s standard deviation was used; DV: random Gaussian noise with a standard deviation of 90% of the feature’s standard deviation was added instead of 120%; DM and DB: a shift of 70% of the feature’s standard deviation was used instead of 90%, and for DP 75% instead of 100%). This analysis assesses whether gamdid’s performance advantages are robust to weaker signals that may be encountered in practice. The results are consistent with the original simulations (figure 6): gamdid maintains conservative FDR control and retains its advantages for shape-based distributional changes (DV, DB), while remaining competitive with distinct for scenarios where location shifts are involved. As expected, the reduced signal does lower the overall power of both methods.

**Figure S5:**
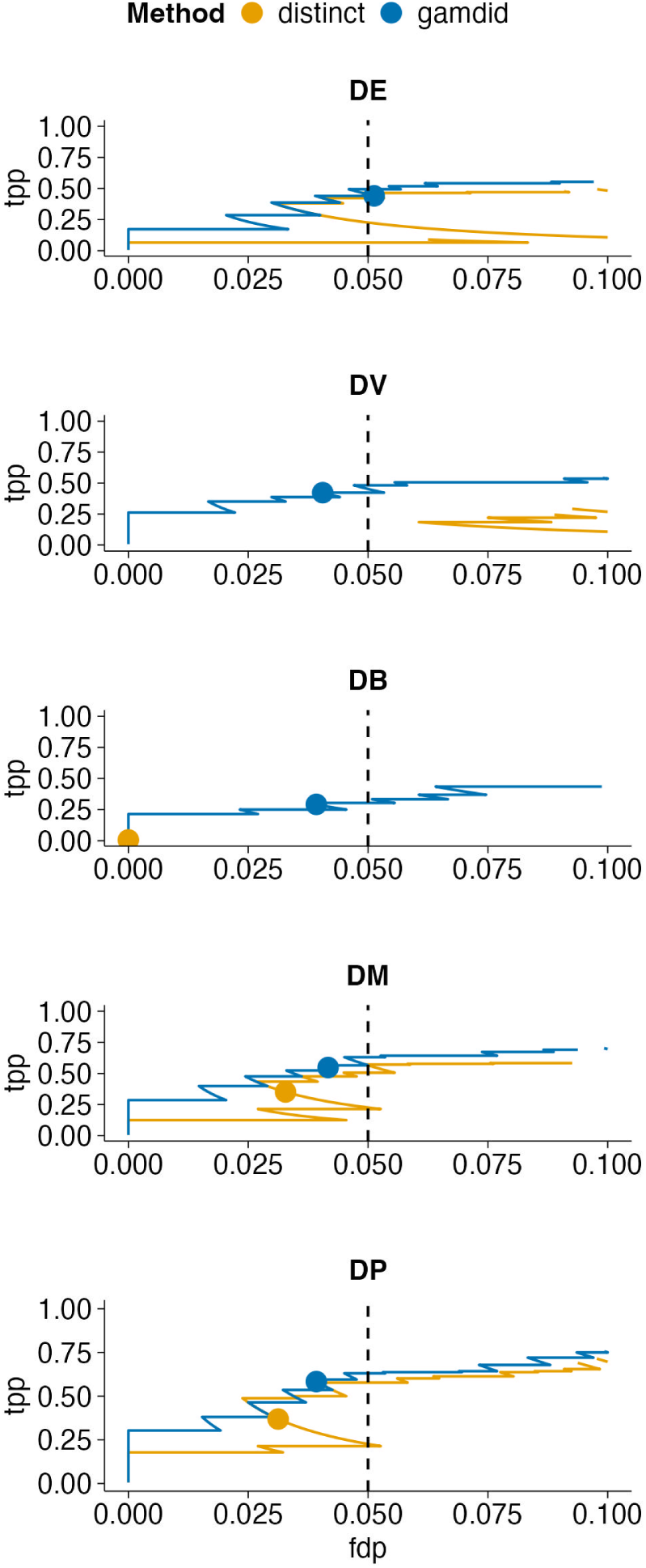
Robustness of gamdid performance under reduced signal magnitude for the Petrosius dataset FDP-TPP curves comparing gamdid and distinct in the Petrosius dataset under simulation scenarios with 25% reduced signal magnitudes, identical to the analysis in figure S4. The results are again consistent with the original simulations (figure 7). The relative improvements over distinct are even more pronounced in the Petrosius dataset than in Leduc, consistent with gamdid’s advantage under smaller sample sizes.

**Figure S6:**
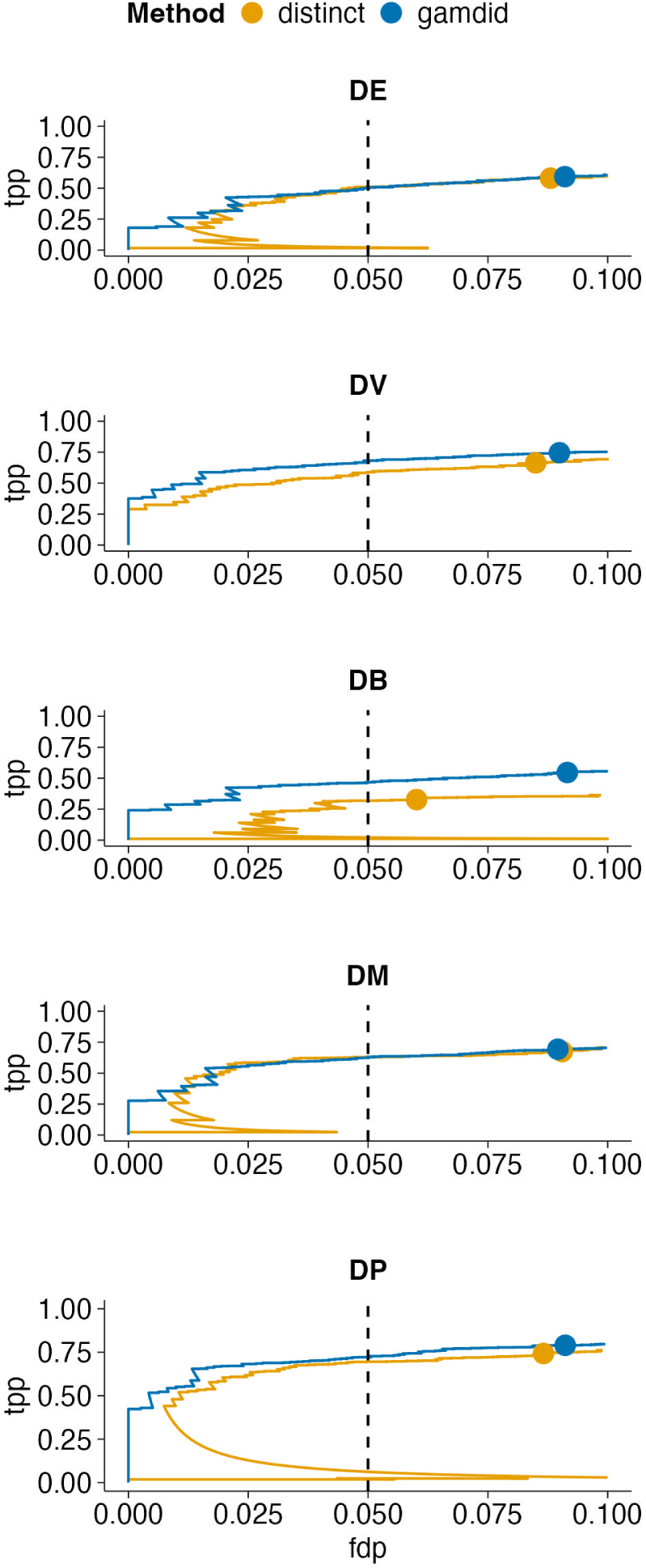
Performance of gamdid and distinct on peptide-level data for the Leduc dataset FDP-TPP curves comparing gamdid and distinct applied to peptide-level assays in the Leduc dataset, rather than the protein-level summarized data used in the main analyses. The results are largely consistent with the protein-level findings (Figure 6). Notably, FDR control is less conservative at the peptide level than at the protein level for both methods. This reflects the well-known challenge in peptide-level data: multiple peptides mapping to the same protein are correlated. Despite this, gamdid provides a flexible and applicable framework across all processing levels of proteomics data.

**Figure S7:**
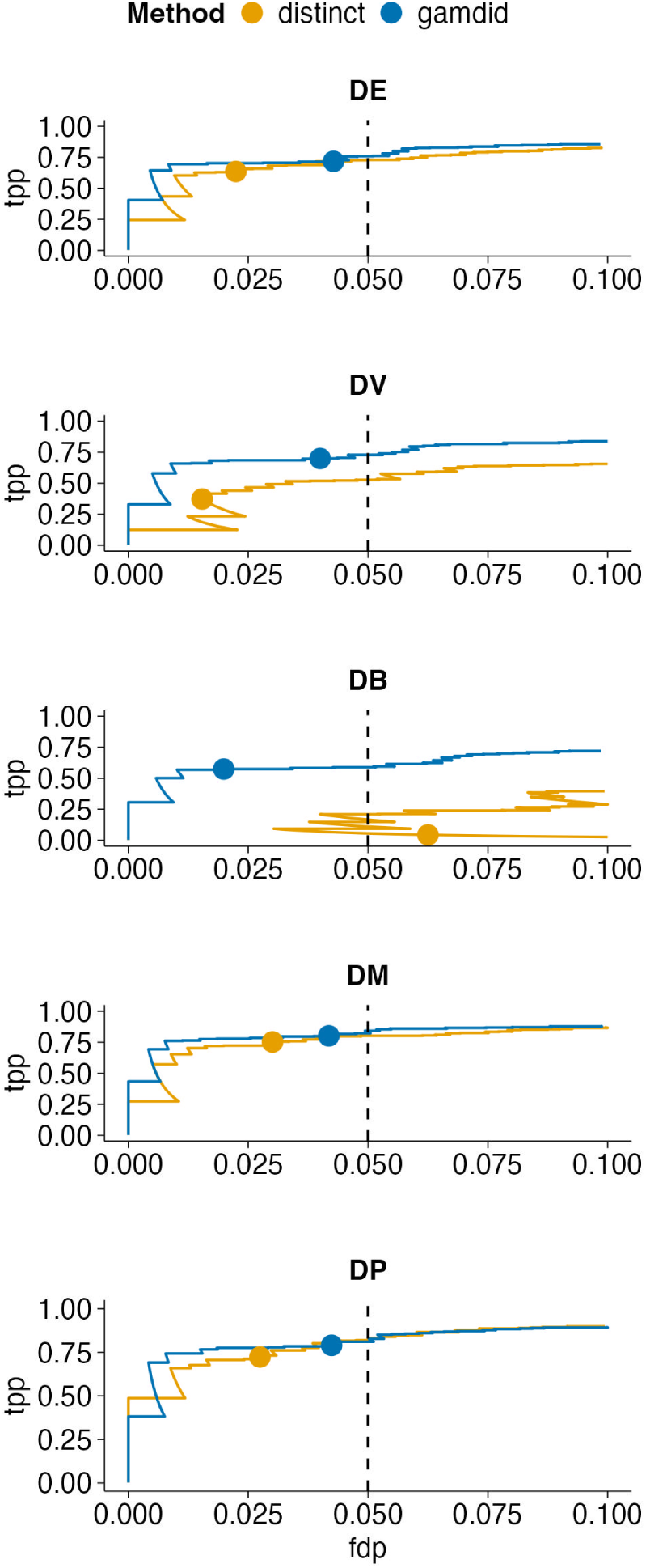
Performance of gamdid and distinct on peptide-level data for the Petrosius dataset FDP-TPP curves comparing gamdid and distinct on peptide-level assays from the Petrosius dataset. The findings are similar to the peptide-level Leduc results (see figure S6): gamdid retains its performance advantages and the less conservative FDR control at the peptide level is again observed for both methods due to inter-peptide correlations.

**Figure S8:**
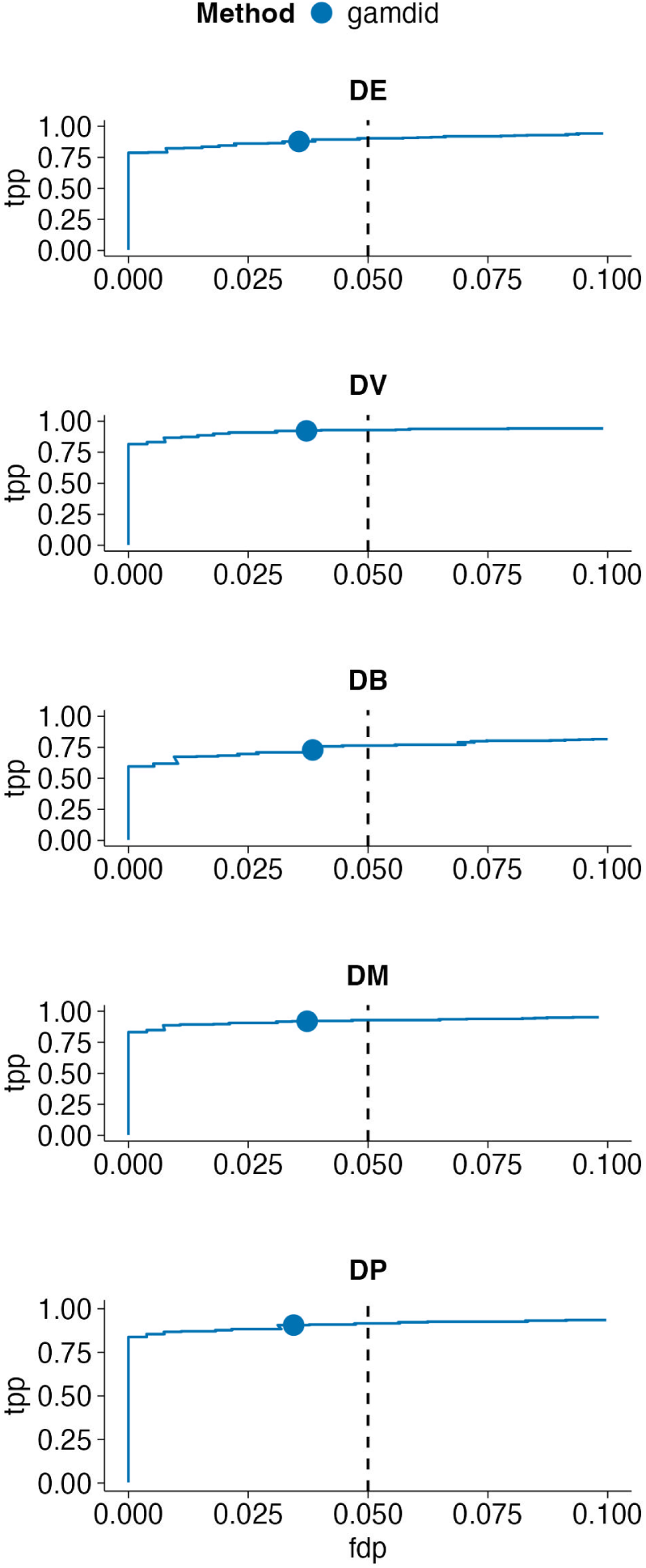
gamdid omnibus test performance in a three-group simulation for the Leduc dataset FDP-TPP curves for gamdid’s omnibus test applied to a three-group comparison in the protein-level Leduc dataset. Cells were randomly assigned to three mock groups (A, B, C), with distributional signal introduced in group B under the same five simulation scenarios as in the two-group analyses. No competing DD method currently supports multi-group omnibus testing, so no benchmark comparison is possible here; these curves demonstrate gamdid’s standalone performance in this setting. Notably, the three-group omnibus test achieves higher power than the corresponding two-group comparisons in figure 6.

**Table S1:** FDR-corrected p-values for features detected by msqrob2 and/or distinct but not by gamdid in the spike-in case study The 8 features that are flagged as significant by msqrob2 and/or distinct but not by gamdid are listed with their FDR-corrected p-values from all three methods. The first 2 features are only detected by distinct. The last 6 features are detected by msqrob2 and missed by gamdid. We see that gamdid consistently returns borderline non-significant p-values for these 6 features. In contrast, distinct produces both relatively high and relatively low p-values for these features. This indicates that only gamdid largely preserves the msqrob2 ranking of DA features.

| msqrob2 | gamdid | distinct |
| --- | --- | --- |
| 0.149 | 0.090 | 0.0454 |
| 0.112 | 0.180 | 0.0454 |
| 0.030 | 0.055 | 0.037 |
| 0.046 | 0.081 | 0.124 |
| 0.048 | 0.081 | 0.045 |
| 0.048 | 0.084 | 0.101 |
| 0.049 | 0.083 | 0.068 |
| 0.049 | 0.086 | 0.124 |

**Table S2:** FDR-corrected p-values for the three features detected exclusively by gamdid in the spike-in case study The three features uniquely identified as significant by gamdid, with FDR-corrected p-values from all three methods. Two of the three features receive very high msqrob2 p-values, confirming that these differences do not involve a detectable mean shift and would be entirely missed by any conventional DA approach.

| msqrob2 | gamdid | distinct |
| --- | --- | --- |
| 0.931 | 0.028 | 0.0516 |
| 0.401 | 0.034 | 0.094 |
| 0.074 | 0.045 | 0.063 |

**Figure S9:**
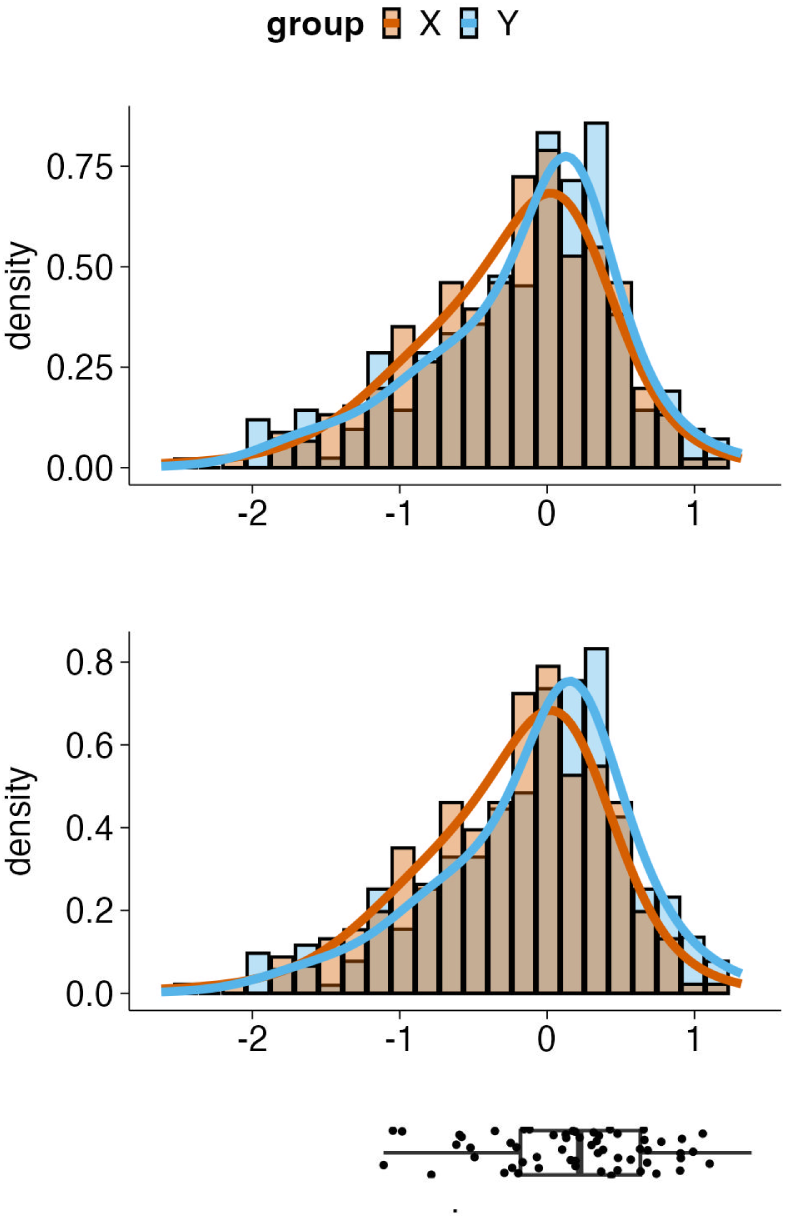
gamdid visualization for a feature detected only by msqrob2 in the spike-in case study (feature 1) The output of gamdid’s visFit function for one of the two features flagged as significant exclusively by msqrob2. **Top**: Mock groups show no distributional difference. **Bottom**: The boxplot below the figures demonstrates where the spike-in melanoma cells are located. After spiking in 10% melanoma cells into group Y, a weak mean-shift appears. No differential region is found by gamdid (no grey shading). This example illustrates a feature type where msqrob2’s dedicated sensitivity for mean shifts gives it a slight edge over gamdid.

**Figure S10:**
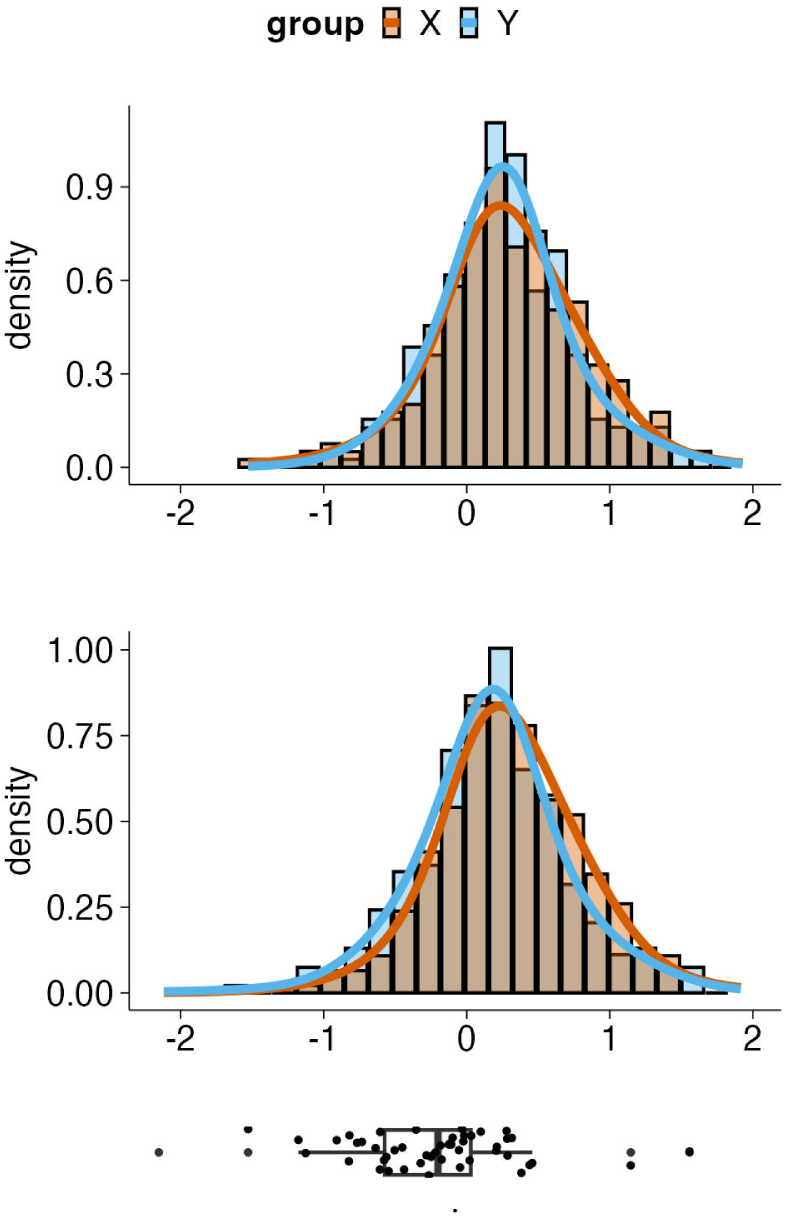
gamdid visualization for a feature detected only by msqrob2 in the spike-in case study (feature 2) The output of gamdid’s visFit function for one of the two features flagged as significant exclusively by msqrob2. **Top**: Mock groups show no distributional difference. **Bottom**: The boxplot below the figures demonstrates where the spike-in melanoma cells are located. After spiking in 10% melanoma cells into group Y, a weak mean-shift appears. No differential region is found by gamdid (no grey shading). This example illustrates a feature type where msqrob2’s dedicated sensitivity for mean shifts gives it a slight edge over gamdid.

**Figure S11:**
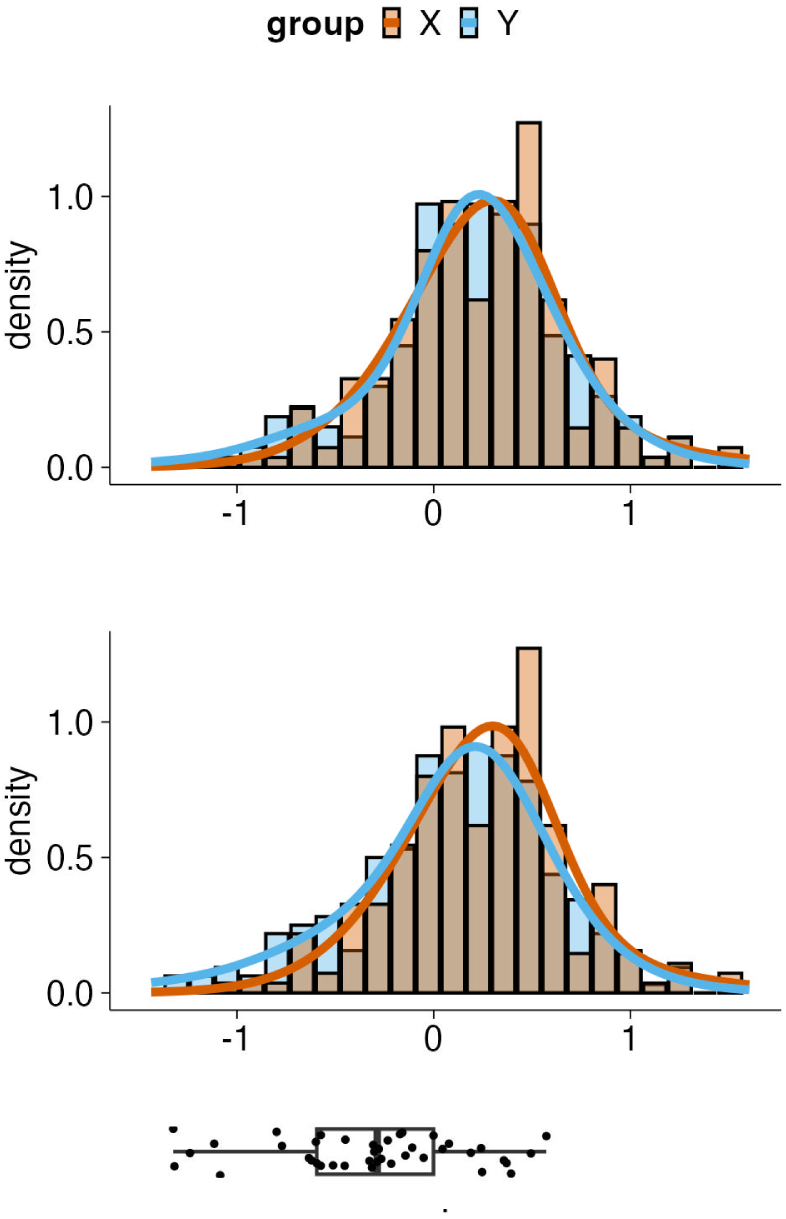
gamdid visualization for a feature detected only by distinct in the spike-in case study (feature 1) The output of gamdid’s visFit function for one of the two features flagged as significant exclusively by distinct. **Top**: Mock groups show no distributional difference. **Bottom**: The boxplot below the figures demonstrates where the spike-in melanoma cells are located. After spiking in 10% melanoma cells into group Y, a weak mean-shift appears. No differential region is found by gamdid (no grey shading). This case illustrates a scenario where distinct’s ECDF-based approach accumulates sufficient evidence to reach significance, while gamdid’s local density approach does not identify a sufficiently concentrated differential region.

**Figure S12:**
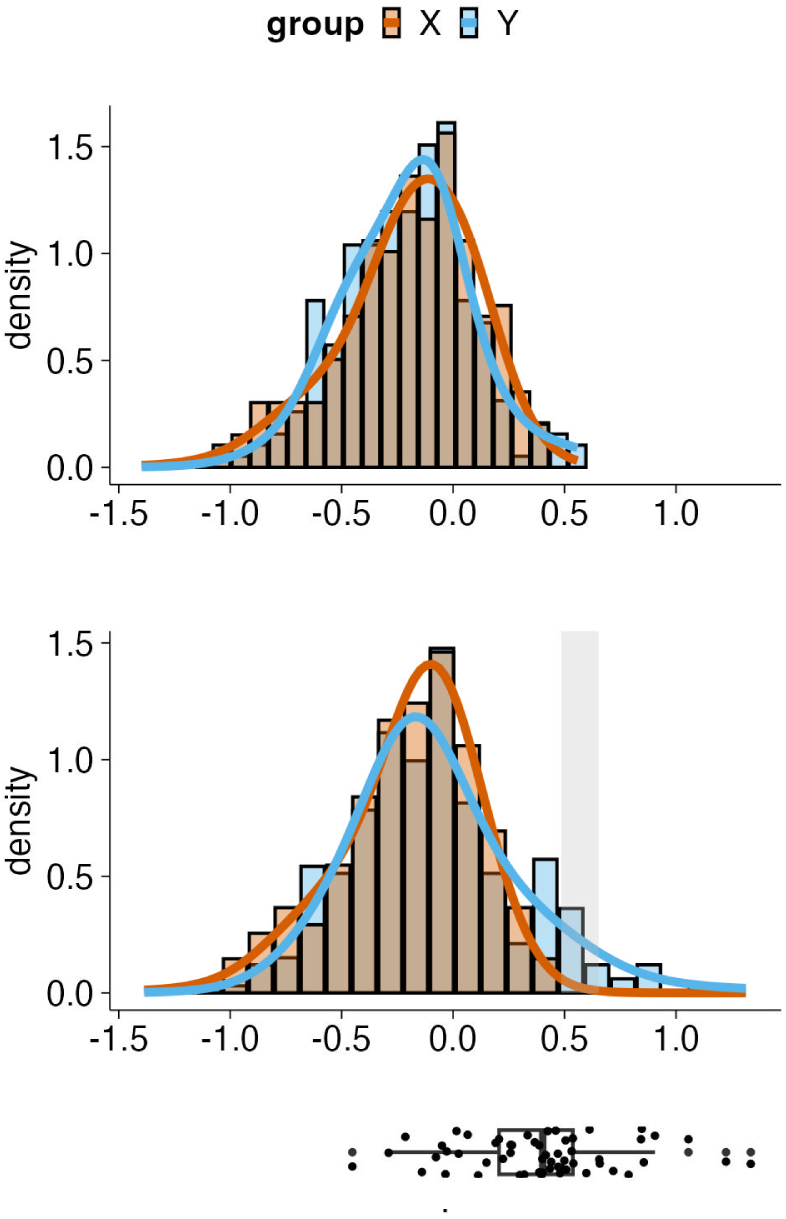
gamdid visualization for a feature detected only by distinct in the spike-in case study (feature 2) The output of gamdid’s visFit function for one of the two features flagged as significant exclusively by distinct. **Top**: Mock groups show no distributional difference. **Bottom**: After spike-in, the density of group Y shifts in the right tail, and gamdid’s difference smoother identifies a differential region (grey shading), despite the FDR-corrected p-value being non-significant (gamdid FDR adjusted p-value = 0.090).

**Figure S13:**
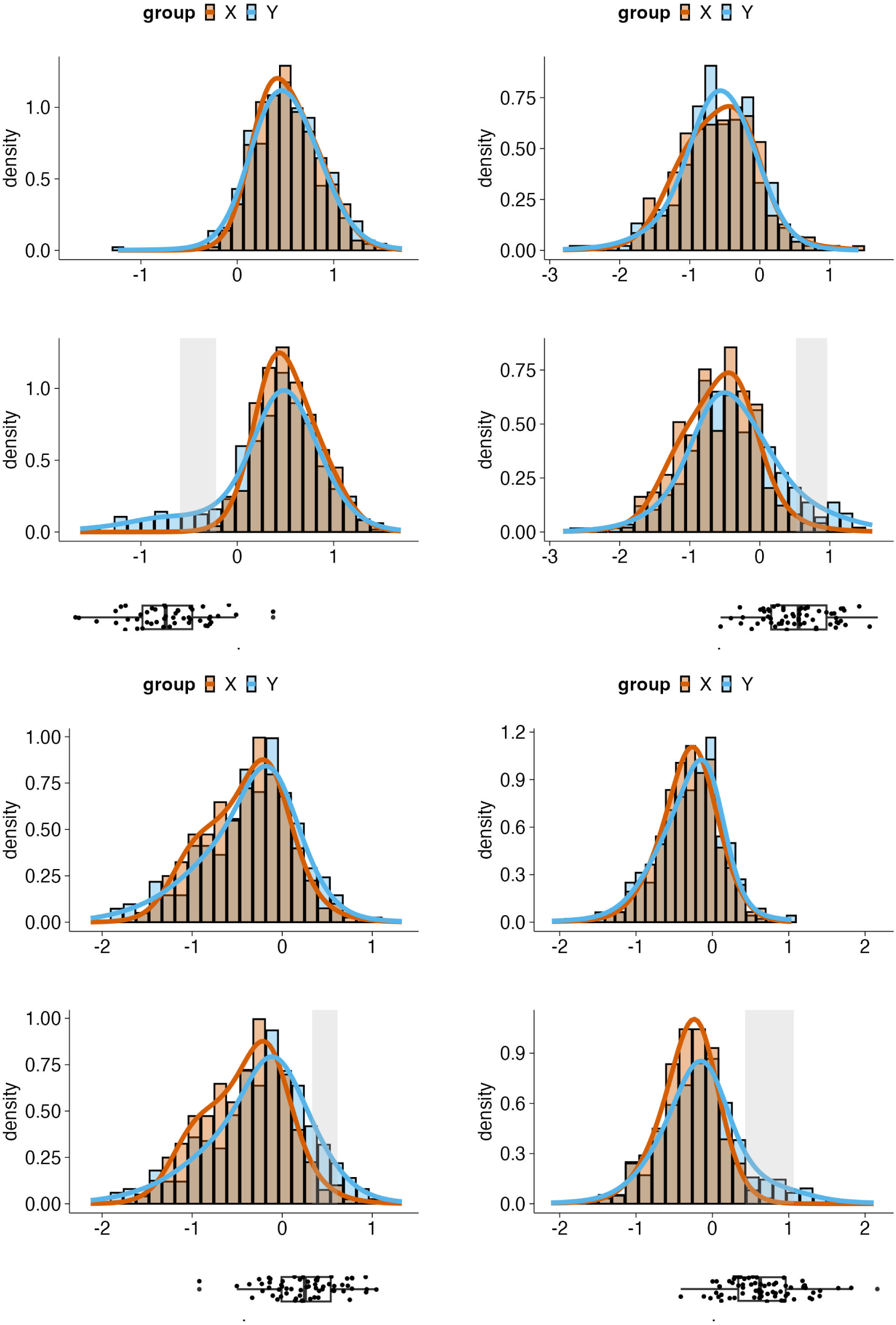
gamdid visualizations for four representative features detected by all three methods in the spike-in case study These four figures follow the same layout as figures S9-S12: mock group comparison (top panel), spike-in comparison (bottom panel), and boxplot of spike-in cell locations. All four features are detected as significant by msqrob2, gamdid, and distinct, forming part of the 54-feature consensus set (see figure 12). In all 4 cases, gamdid’s visualization reveals that the distributional difference extends beyond a simple mean shift: these features are DD due to the subpopulation of spike-in cells rather than DA. Moreover, the spiked-in melanoma cells align closely with the detected differential region (grey shading) in each example. This demonstrates that even within the set of features identified by conventional DA methods, gamdid provides additional interpretive value by characterizing how and where the distributions differ. This information is not accessible from the fold change and p-value from a conventional DA method. These examples underscore the broader argument that accounting for full distributional differences, rather than mean shifts alone, leads to richer biological insights.

## Generalized likelihood ratio test (GLRT)

The null model (equation 11) can be compared to the full model (equation 9) using a GLRT (see equation 16).

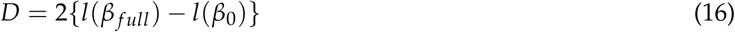

where *l*(*β f ull* ) and *l*(*β*_0_) are the maximized log-likelihoods for the full and the null models, respectively. Under *H*_0_, *D* asymptotically follows a chi-squared distribution (see equation 17).

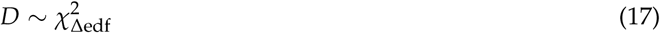

where Δedf = edf *_f ull_* − edf_0_. edf *_f ull_* and edf_0_ are derived from equation 8. The p-value is then 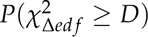 [26]

This approach requires fitting two models and does not directly provide pairwise comparisons if more than two groups exist.

## Notes

### Competing Interest Statement

The authors have declared no competing interest.

### Funding

- Ghent University

